# Reward prediction-errors weighted by cue salience produces addictive behaviors in simulations, with asymmetrical learning and steeper delay discounting

**DOI:** 10.1101/2023.03.19.533364

**Authors:** Shivam Kalhan, Marta I. Garrido, Robert Hester, A. David Redish

## Abstract

Dysfunction in learning and motivational systems are thought to contribute to addictive behaviours. Previous models have suggested that dopaminergic roles in learning and motivation could produce addictive behaviours through pharmacological manipulations that provide excess dopaminergic signalling towards these learning and motivational systems. Redish 2004 suggested a role based on dopaminergic signals of value prediction error, while Zhang et al. 2009 suggested a role based on dopaminergic signals of motivation. Both these models present significant limitations. They do not explain the reduced sensitivity to drug-related costs/negative consequences, the increased impulsivity generally found in people with a substance use disorder, craving behaviours, and non-pharmacological dependence, all of which are key hallmarks of addictive behaviours. Here, we propose a novel mathematical definition of salience, that combines aspects of dopamine’s role in both, learning and motivation, within the reinforcement learning framework. Using a single parameter regime, we simulated addictive behaviours that the Zhang et al. 2009 and Redish 2004 models also produce but we went further in simulating the downweighting of drug-related negative prediction-errors, steeper delay discounting of drug rewards, craving behaviours and aspects of behavioural/non-pharmacological addictions. The current salience model builds on our recently proposed conceptual theory that salience modulates internal representation updating and may contribute to addictive behaviours by producing misaligned internal representations (Kalhan et al., 2021). Critically, our current mathematical model of salience argues that the seemingly disparate learning and motivational aspects of dopaminergic functioning may interact through a salience mechanism that modulates internal representation updating.

## Introduction

The ability to adapt behavior in response to the ever-changing environmental contingencies is a central component of successful decision-making. Intelligent behavior entails selecting the best actions given the information available and the biological constraints on computation in the brain. In a world with an abundance of sensory information, animals learn from and use information based on a number of key factors, including how well that information predicts the future. The Rescorla-Wagner (RW) model of learning is one way this type of cue-outcome associative learning can be computed (Rescorla & Wagner, 1972). The central component of the RW model is that learning only occurs when there is a prediction-error generated, which occurs when expectations of reward (or lack thereof) are violated. The RW model is a simple, yet powerful, way of explaining several aspects of learning. However, it has limitations. One limitation is that it cannot explain second or higher order conditioning, which is where an associative relationship is formed between the second order cue that predicts the first order cue which predicts the outcome. Given that many real life forms of learning involve second or higher orders of conditioning (e.g., money is a second order predictor of many outcomes like food and shelter), this is an important limitation to overcome. Another limitation in the RW model is that each trial is treated as a discrete temporal object. In reality, a trial is one part of a continuous sequence of events, including the *temporal* relationship between the cue and the outcome, which is one major determinant of learning. Temporal difference reinforcement learning (TDRL) models extend the RW model and overcome these two limitations.

The TDRL model proposed that agents (animal, robots, or simulations) select actions to maximize future reward and minimize future costs (Sutton & Barto, 1998). These models usually learn a value function over a finite Markovian decision process (MDP) where certain *observations* are expected to occur in a specific *state*. These observations could be a cue, a reward, or a cost (negative reward). In these models, value is defined as the discounted, expected future reward and a value is associated with each state. Based on its observations which predict the state the agent is in, agents can select actions that lead to the states with the highest values (high reward, low cost). In these models, agents learn and select actions that increase the likelihood of entering states with high value while also avoiding entering states with low values. Computing these distinct states and values through MDPs allows a simple, yet powerful, way of understanding decision-making processes as well as how and why they may go awry (i.e., when animals choose states that do not appear to have high values for them and are maladaptive). We specifically define an internal representation here as a *state-space* model within the TDRL framework that the agent can use to select actions based on value computations.

The value of a given state, at time t, can be denoted as *V(S_t_)* and is defined as the sum of expected future rewards, discounted by the delay (0 < γ < 1) to the reward:

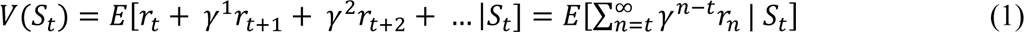

According to this value definition, the further in time the rewards are, the more their values are discounted. Importantly, from this definition it directly follows that the value of a given state at time t is equal to the immediate reward received in that state plus the discounted value of the next state, at time t+1:

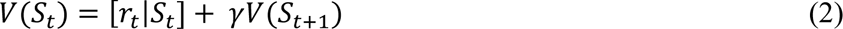

If the two sides of the above equation are not equal, a prediction-error (δ) is generated, which is the difference between the two sides of the equation, where the outcome ([*r_t_*|*S_t_*] + γ*V(S_t+1_)*) is different from the expected value, (*V(S_t_)*). Therefore, the *V(S_t_)* needs to be updated/learnt to better reflect the outcome. Hence, equation 2 is simply rearranged to the following, capturing the prediction-error:

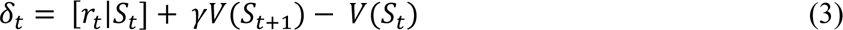

This prediction-error is used to learn and update the expectation, where the old expectation (*V(S_t_)old*) is updated to the new expectation (*V(S_t_)new*) as below:

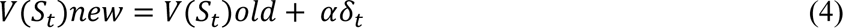

where α is the learning rate, a number between 0 and 1. Importantly, within this framework, if the prediction-error is 0, there is no learning/updating of the state value and the new value is identical to the old value. However, if there is a positive prediction-error (i.e., the outcome was better than expected), the value of the state is increased. Therefore, actions that lead to that high value state will be reinforced. But, if the prediction-error is negative, that is, the outcome is worse than expected, then the actions leading to this low value state are downregulated. In this way, prediction-errors can be useful ways of constantly learning and updating cue-outcome relationships, by reinforcing actions that lead to states with high values. The main point here is that positive prediction-errors reinforce actions through value increase, but negative prediction-errors downregulate actions through value decrease.

Since learning in these models depends entirely on prediction-error, both the RW and the TDRL models, cannot account for the concept of *latent inhibition.* This is where pre-exposure to cue A (with no outcome) in phase 1, delays the later learning of the relationship where cue A predicts outcome in phase 2 (Lubow & Moore, 1959). The RW model incorrectly predicts that pre-exposure to cue A should not impact learning in phase 2. Latent learning (i.e., learning without an explicit outcome) is a key component in optimizing behavior (Tolman, 1948), and that the RW model cannot explain this, is a significant limitation. To address this limitation, the concept of *cue salience* was added to the RW model (Mackintosh, 1975; Pearce & Hall, 1980). The cue salience models argued that the pre-exposure reduced the salience of the cue because it had no value in predicting the outcome in phase 1, and so it should take longer to learn as the salience needs to be increased after the repeated pairings of cue A and outcome in phase 2. At its core, the cue salience models suggested an attentional weighting mechanism for cues depending on the salience, which one could expect to also influence learning/updating rather than just the expectation of the outcome. The larger the salience, the greater updating of the internal representation, i.e., learning. We recently proposed a conceptual theory suggesting that aberrations in these cue salience/attention weighting mechanisms could play a role in explaining some learning and decision-making aspects in addictive-like behaviors (Kalhan et al., 2021). We proposed that a consequence of drug cues gaining a high salience is asymmetric learning, where a misaligned internal representation is formed, and this misaligned internal representation is then used to produce maladaptive drug-related behaviors. Using this concept, the primary aim of the present paper is to add a cue salience weighting mechanism to the TDRL model, conceptually similar to how a salience factor was added to the RW model (Mackintosh, 1975; Pearce & Hall, 1980), such that learning is also influenced by the cue itself and not only the outcome. In so doing, we propose a neural computation that produces key aspects of addictive behaviors.

The Redish (2004) model attributed addictive behaviors to reinforcement learning computations gone awry. It combined two key concepts. The first concept was built on the hypothesis that the prediction-error term is encoded within dopaminergic cell firing in the ventral tegmental area (Schultz et al., 1997). The second key concept was that many drugs of abuse release dopamine, either directly or indirectly (Nutt et al., 2015; Ritz et al., 1987). Therefore, Redish (2004) proposed that actions that lead to drug-receiving states produce a positive prediction-error on drug receipt due to an increase in dopaminergic activity from the drug, which cannot be compensated for, under normal reinforcement learning processes. As a result, the value of these drug states continues to increase, reinforcing both drug-seeking and drug-taking actions. Redish (2004) proposed a modified prediction-error equation based on the drug’s non-compensable dopamine release (ncDA):

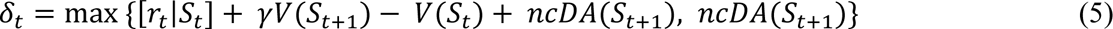

Using this prediction-error equation, Redish (2004) demonstrated via simulations that agents built based on this equation are less likely to choose a non-drug option over a drug option of the same reward magnitude to reduce drug-seeking actions, the non-drug rewards need to be of greater magnitude than the drug rewards. In this model, the more times the agent took the drug action, the greater the alternative non-drug reward needed to be to decrease drug-seeking actions. Additionally, agents demonstrated inelasticity in that they increasingly became less sensitive to costs the more drug actions they took. Overall, these simulations suggested that the non-compensable dopamine release by drugs, causing constant positive prediction-errors, may be one possible way in which drug-seeking actions are continually reinforced in some people with substance use disorders.

Another influential account of dopamine’s involvement in addiction is the *incentive salience* account (Robinson & Berridge, 1993). According to that account, dopamine is a driver of addictive behaviors through its role in triggering motivational ‘wanting’, and less so due to its role in learning. The key idea here is that repeated exposure to drugs causes dopaminergic circuits that attribute salience towards drug predictive cues to become hypersensitized (Robinson & Kolb, 2004; Singer et al., 2009; Steketee & Kalivas, 2011). As a result, drug predictive cues have a high salience value when encountered, triggering a strong motivational ‘wanting’ of drugs and increasing the persistence in drug-seeking actions. More recently, (Zhang, Berridge, Tindell, Smith, & Aldridge, 2009) proposed a neurocomputational account for this theory. The authors proposed that when a drug cue with high salience is paired with a given reward, the high salience makes that reward more reinforcing than it normally would. Mathematically, the authors proposed the following equation:

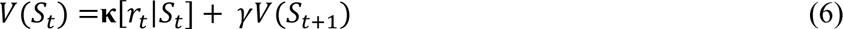

where, **κ** is a salience factor that can dynamically modulate the value computation of a given state. Here, a higher salience would mean a greater value increase towards that state, and in this way, agents persist in drug-seeking behaviours.

Critically, both the Redish (2004) and the Zhang et al., (2009) models suggest that drug states have higher values that lead to the persistence in drug-seeking behaviours. The Redish (2004) model suggests that the value increase is incremental, through prediction-error induced learning. The Zhang et al., (2009) model has an added salience factor which can be dynamically modulated through physiological changes, and can thus account for fast, dynamic fluctuations in behaviour, without incremental relearning.

Both models present limitations. Critically, the Zhang et al., (2009) model does not mathematically define the salience factor, and how it may be generated within the model itself. Therefore, the agents simply behave as if reward is increased multiplicatively. For example, if the salience is 2, and reward is 5, the agent will act identically to if the reward was 10. However, addictive drugs are not necessarily more ‘rewarding’. In fact, they come with great negative consequences (i.e., poor health). Second, the model does not account for negative prediction-errors or drug related costs. For example, if the drug cue, which has a high salience, is followed by an unexpected cost (negative reward prediction-error), it would make the cost greater than it is. This should greatly decrease the value of drug states, and the agent should be much less likely to persist in drug-related behaviours compared to any other behaviour with a lower salience. Of course, people with substance use disorders often find drug-seeking actions difficult to decrease, and many are compulsive in their behaviours, where drug states are less sensitive to devaluation (Everitt & Robbins, 2005, 2013; Lüscher et al., 2020).

A limitation of the Redish (2004) model is that it cannot account for the cue-triggered dynamic fluctuations in behaviour, because any changes would have to be from the incremental relearning of value. Furthermore, equation (5) as written does not allow delta to be negative, even for non-drug rewards – if ncDA=0 then the minimum delta would be 0. Later studies by that laboratory proposed that extinction (lack of delivered reward) occurs via a different process (Redish et al 2007), but dopamine does decrease in real paradigms (Abraham et al., 2014; Schultz et al., 1997), so processes need to be in place for fluctuations in dopamine in both positive and negative prediction-errors, which will thus require a modification of equation (5). Further, neither model has addressed the increased impulsivity, seen as a specifically steeper delay discounting of drug rewards. This steeper delay discounting is a hallmark deficit consistently found in people with substance use disorders and is even used as a biomarker for addictive behaviours (Bickel et al., 2014; Madden & Bickel, 2010). Additionally, the two models do not account for craving behaviours, which is also not commonly accounted for in many models of addiction (but see a recent review by Mollick & Kober (2020)).

To address these limitations, we propose a new neurocomputational model for addictive-like behaviours, which is a partial combination of the two models, with a novel mathematical definition of salience. Specifically, we aimed to produce a model where 1) salience is mathematically defined within the reinforcement learning framework, 2) the salience factor can account for sudden cue-triggered fluctuations in behaviours, 3) negative prediction-errors are also accounted for within the model, 4) the hallmark deficit of steeper delay discounting for drug rewards is accounted for, and 5) the model can also explain some craving behaviours, all of which can be induced to the agents mathematically within a single parameter regime.

## METHODS

We start from a modification to the Redish (2004) model when calculating the prediction-error, where if the agent received a non-drug reward (ncDA = 0), the normal TDRL formula would be used:

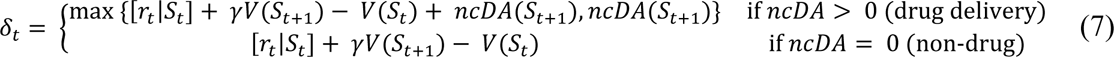

Because of this modification, less-rewarding-than-expected non-drug rewards can now provide negative prediction-errors, which was not possible under the original Redish (2004) formula.

We then mathematically define salience, κ, as:

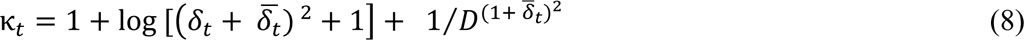

where, D is the delay to the reward. In the first component of the equation 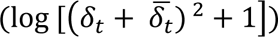, a high average prediction-error (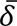) state will place more salience on the positive prediction-errors, and at the same time a lower salience on negative prediction-errors. The second component 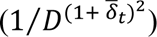 models the role of delay discounting in salience as salience being inversely proportional to time/delay. Additionally, the greater the average prediction-error (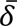), the lower the salience placed on the delay. We then use this salience factor to update the internal representation multiplicatively:

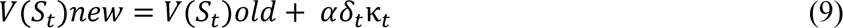

Here, value updates are proportional to the salience, and importantly, salience is a *modulator* of these internal representation updates, hence, if the prediction-error is 0, there is no effect of salience.

World states were implemented as Markovian processes wherein an agent transitions from one state to the next at time t+1, with a given transition probability matrix (Sutton & Barto, 1998). Importantly, in all simulations, state 1 is an inter-trial interval state (ITI) which is made up of many different states that the agent can go back to (at random) after having completed a trial. Because all simulations used a micro-agent model (Kurth-Nelson & Redish, 2009; Redish, 2004) where one agent (macro-agent) consisted of 100 micro-agents that differ in their discounting factor (**γi**), prediction error is distributed across that ITI, slowing down cyclical learning (Kurth-Nelson & Redish, 2009; Redish, 2004). Discounting factors **γi** were randomly assigned to each micro-agent from a uniform distribution with factors between 0 and 1. Each micro-agent computed its own prediction-errors and state value updates based on their discounting factor, with the decisions made by the macro-agent based on these individual value updates (benefit; B, eq 10) using the softmax function (eq 11):

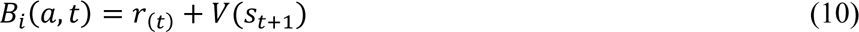

Where, *B_i_(a, t)* is the benefit of taking action *a* at time t, for micro-agent *i*, *r_(t)_* is reward at time t and *V(s_t+1_)* is the value of the next state (the state reached at time t+1), given action *a* is taken. Each micro-agent performs this benefit calculation individually, which are then averaged across all the micro-agents, and this average is taken as the averaged benefit of taking action *a* by the macro-agent, 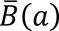. Decisions were made by the macro-agent. The probability of taking each action is computed using the softmax action selection function:

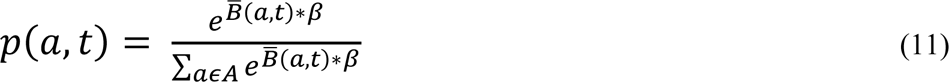

Where, *p(a, t)* is the probability of taking action *a* at time t, and β is the inverse temperature (where β = 0 provides random responding and β = 1 always takes the action with the greatest benefit).

Finally, we repeated each simulation 10 times, each with a different set of 100 micro-agents to capture any variability due to differences in the average discounting factor. The parameters of the learning rate (α) used was 0.05 and inverse temperature (β) was 0.5, in all simulations, unless stated otherwise. All simulations were done using MATLAB-2017b, with figures generated using RStudio (version 4.1.2). See supplementary materials for additional details of each simulation.

### Justification of modelling choices

#### Modelling asymmetrical learning from positive and negative prediction-errors

The definition of salience proposed here, particularly the first component 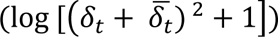, builds on the findings of Frank et al., (2004) showing that when people with Parkinson’s disease were on dopaminergic medication (i.e., they were in a high dopamine state), they learned more from positive feedback than from negative feedback. This phenomenon is very similar to the learning behaviours generally observed in people with substance use disorders being more sensitive to the positive outcomes and less sensitive to negative outcomes. Given that most drugs of abuse release dopamine (Nutt et al., 2015), a consequence of drug taking may be to drive the agent into a high dopaminergic state (high 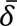). Linking this with the Redish (2004) model, a consequence of the ncDA release from drugs is a constant positive prediction-error. Therefore, we added a factor which is calculated based on the prediction-error, and that is the average prediction error (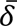). The higher the average prediction-error, the higher the dopaminergic state that the agent is in, caused by the ncDA release from the drug. Therefore, a high average prediction-error, encoding a high dopamine state, will also attribute increased salience to positive prediction-errors and at the same time, attribute less salience to negative prediction-errors, as in equation (8). As a result of this asymmetric salience attribution, there will be less learning/updating from negative prediction-errors, but more so from positive prediction-errors. Therefore, using our salience formula, we can mathematically capture the phenomenon observed by Frank et al., (2004) where there is asymmetric learning from positive and negative feedback depending on the dopaminergic state.

#### Modelling steeper discounting for drug rewards

The next aim was to account for the steeper delay discounting for drug rewards found in people with substance use disorder, which speaks to the second component of equation 8 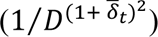. The *temporal construal* theory by Trope et al., (2003) suggested that things further in time are discounted because they have a reduced concreteness, where the future has a higher level construal and is represented more abstractly. The future also has increased uncertainty. Accounting for these factors produces increased discounting of reward values as time to reward increases (Madden & Bickel, 2010). Equation 7 accounts for this phenomenon through the discounting factor (γ), which reduces the value of rewards that are further away in time. Interestingly, when the future is made more precise and is concretely represented via episodic future thinking, by reminding participants what they will be doing that day in the future, discounting is less steep (Peters & Büchel, 2010). Koffarnus et al., (2013) suggested that this may occur due to an increased salience in the future event representation. Therefore, one interpretation of why future rewards are discounted more is that the future has less salience than the present, suggesting that salience is inversely proportional to the delay (D), which we thus added as a component in our salience formula. However, we wanted to link this with addiction and the increased dopamine state caused by the high average prediction error (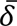). Pine et al., (2010) found that when human participants were given L-dopa, which increases dopaminergic levels, they discounted faster compared to when they were given a placebo drug. The Niv et al., (2007) theory of opportunity costs suggested via simulations that the greater the average reward rate for the animal (which is when there is a high average prediction error and dopaminergic state), the greater the vigor with which the animal responds to opportunities. This idea has since been demonstrated experimentally in humans (Beierholm et al., 2013). The reasoning for this positive correlation between reward rate and vigor was suggested to arise because the cost of not acting (or of acting slowly, i.e., with less vigor) is higher in a reward-rich environment. This suggests that discounting may also be steeper for delayed rewards when the animal is in a rich environment. Further, Ballard et al., (2015) found that in people with a methamphetamine dependence, the lower the dopamine receptor D2/3 availability, the steeper the rate of discounting. Overall, these accounts suggested that the higher the dopamine levels, the steeper the discounting, and therefore the high average prediction-error produced steeper discounting in our model, through modulating the salience placed on the delay. The higher the average prediction-error, the less the salience is placed on the delay, and the steeper the discounting. Importantly, this steeper discounting effect for drug rewards was a *consequence* of the model as opposed to being built into the model, as we did not directly manipulate the discounting factor (γ) to produce this effect.

#### Modelling hyperbolic instead of exponential delay discounting of drug and non-drug rewards

The primary motivation for using the model with micro-agents instead of a more traditional reinforcement learning model (with one agent) was to produce hyperbolic delay discounting of rewards (instead of exponential) which is closer to how biological agents discount delayed rewards (Madden & Bickel, 2010). The basic reinforcement learning equation (Eq. 7) discounts delayed rewards exponentially, however, a distribution of these exponentially discounting agents produces hyperbolic discounting (Kurth-Nelson & Redish, 2009). Moreover, a distribution of differentially weighted prediction-errors are likely encoded in a distribution of dopaminergic neurons that encode a reward prediction-error (Dabney et al., 2020), with a distribution of discounting factors possibly encoded in a gradient-like manner within the striatum (Onoda et al., 2011; Tanaka et al., 2004; W. Wei et al., 2021).

### Methods Summary

In the results section, we will compare how the Redish 2004 model, the Zhang et al. 2009 model, and our current model react to various state space contingencies under drug and non-drug rewards. We will refer to our current model as a Salience Misattribution Model for Addiction (SMMA). Table 1 shows the equations used for each model.

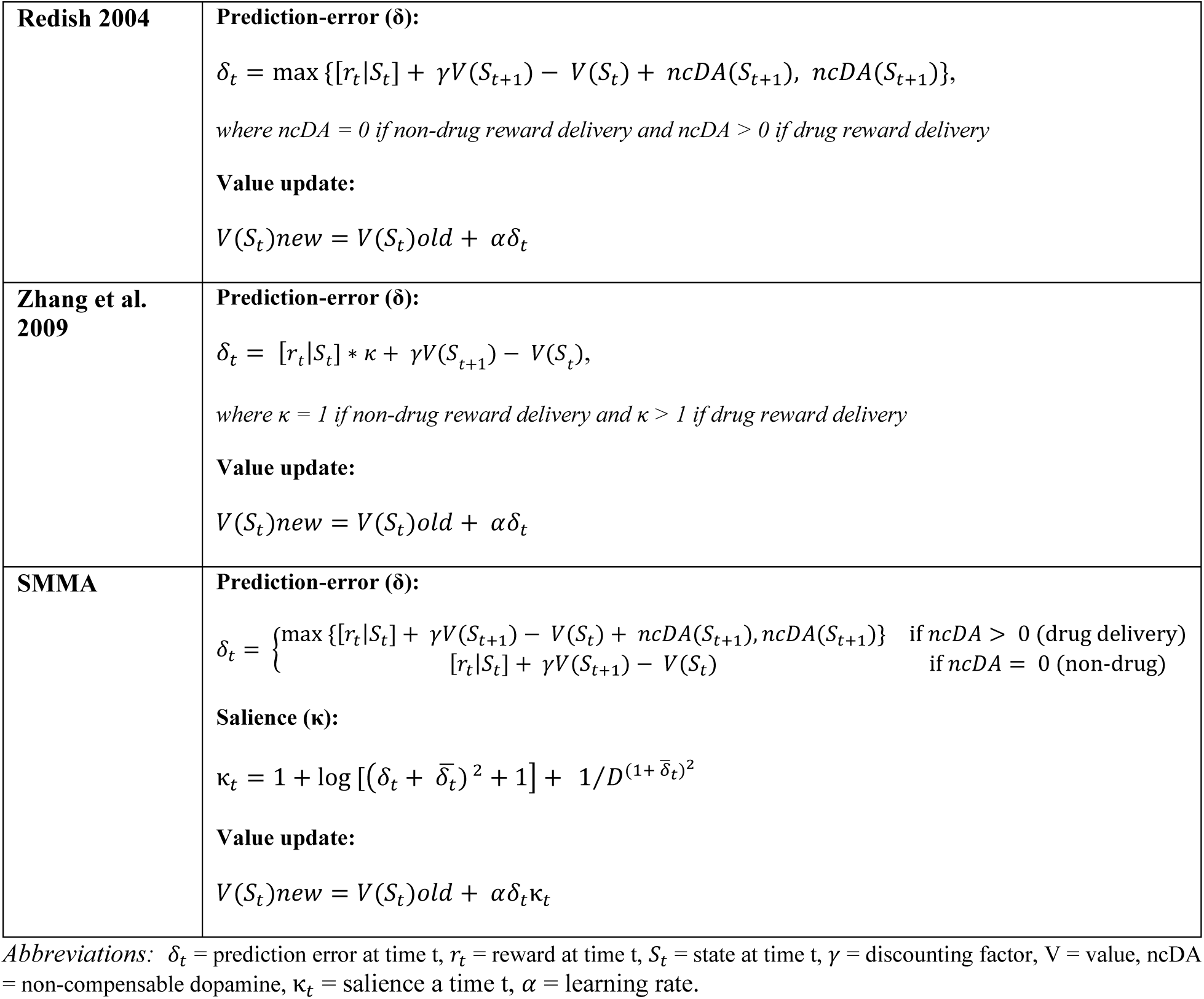

## RESULTS

### Increased salience on drug cues

Owing to a higher salience, drug cues generally show larger reinforcing effects on behaviour than cues leading to non-drug rewards (Adinoff, 2004; Carter & Tiffany, 1999; Lubman et al., 2000, 2007, 2008, 2009). In reinforcement learning models, these effects are hypothesized to arise from the differences in their effects on value updates and prediction-errors. The Redish 2004 model does not have a salience factor. However, it produces unbounded value increase towards drug states (Figure 1e) due to the non-compensable positive prediction-errors that drugs create (Figure 1f). The Zhang et al. 2009 model does have a salience factor, and, increasing salience, without changing the delta equation, accelerates learning – but does not change the value plateauing or prediction-error reaching zero (Figure 1h and 1i). The SMMA model, however, includes both of these effects: accelerated learning as the salience increases, and an always-positive prediction-error, which causes the value update to increase without bound (Figure 1k-o). In the SMMA model, these effects interact - the salience accelerates the unbounded value update, due to being multiplied by the prediction-error and causing a greater value update, with a greater salience.

**Figure 1.**
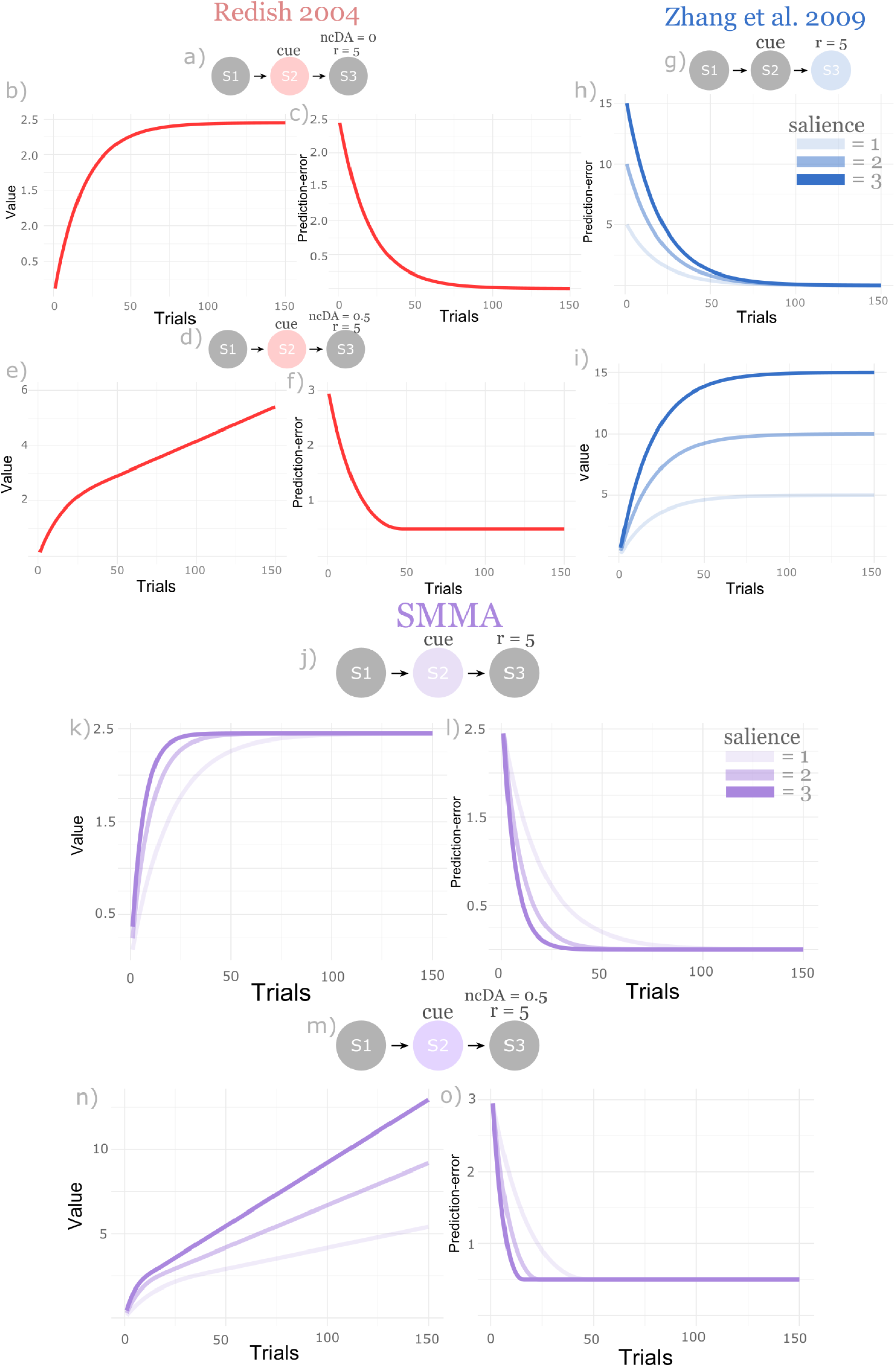
Drug and non-drug reward value and prediction-errors based on the Redish 2004, Zhang et al. 2009 and the SMMA models. **a)** The state space for the Redish 2004 model, for non-drug rewards where ncDA = 0 and r = 5 at state 3 (S3). The value and delta from the cue state, state 2 (S2), is plotted using this model. **b)** value (y-axis) and the number of trials (x-axis) for non-drug rewards using the Redish 2004 model. **c)** the non-drug reward prediction-error (y-axis) and the number of trials (x-axis). **d)** the state space for the Redish 2004 model, for drug rewards where ncDA = 0.5 and r = 5 at S3. **e)** unbounded value increase from drug rewards due to ncDA > 0, caused by the **f)** constant positive prediction-errors from drug rewards. **g)** The state space for the Zhang et al. 2009 model, with cue salience from S2 multiplied by the reward at S3. **h)** as the salience increases, so does the magnitude of the prediction-errors, but it eventually reaches 0. **i)** as the salience increases, the value also increases, but plateaus (e.g., if r = 5, and salience = 2, value will plateau at 10). **j)** The state space used for the SMMA model, for non-drug rewards where ncDA = 0 and r = 5 at S3. The value and delta from the cue state, state 2 (S2), is plotted using this model. **k)** as salience increases value increase accelerates and **l)** prediction-errors reduces faster, but both plateau at the same point, irrespective of the salience. **m)** the state space for the SMMA model, for drug rewards where ncDA = 0.5 and r = 5 at S3. **n)** value increases without bound for drug rewards, and increasing salience accelerates this value increase. **o)** the increase in salience accelerates prediction-error learning but plateaus at the same point (above 0), irrespective of salience.

### Lever presses for drug rewards

Reward predicting cues, including drug cues, generally have high salience, and as a result reinforce behaviours that lead to them (e.g., lever presses for a reward following the presentation of a reward predicting cue) (Domjan, 2014; Flagel et al., 2009; Uslaner et al., 2006). Here we simulated this effect of cue salience on pressing a lever for a drug or a non-drug reward using the Zhang et al. 2009 model, Redish 2004 model, and the current SMMA model (Figure 2). A key component of the Zhang et al. 2009 model was that as salience increases for drug cues, so does the proportion of drug actions, and that this is dynamic, without any further learning. The Redish 2004 model does not have a salience factor, and any increases in drug behaviours using this model is through the incremental unbounded value increase by changing the delta equation, not through any changes in salience, which is why there is a flat line produced when simulating the effect of changing salience using the Redish 2004 model (Figure 2; red line). However, the SMMA model is able to simulate the increased lever presses for drugs through changes in the salience factor itself, just as is the case in the Zhang et al. 2009 model. In both simulations, using the Zhang et al. 2009 and the SMMA model, all parameters, except the salience factor, remained the same. Therefore, the increase in lever presses is caused by the dynamic salience factor itself, and not changes in reward value for the drug.

**Figure 2.**
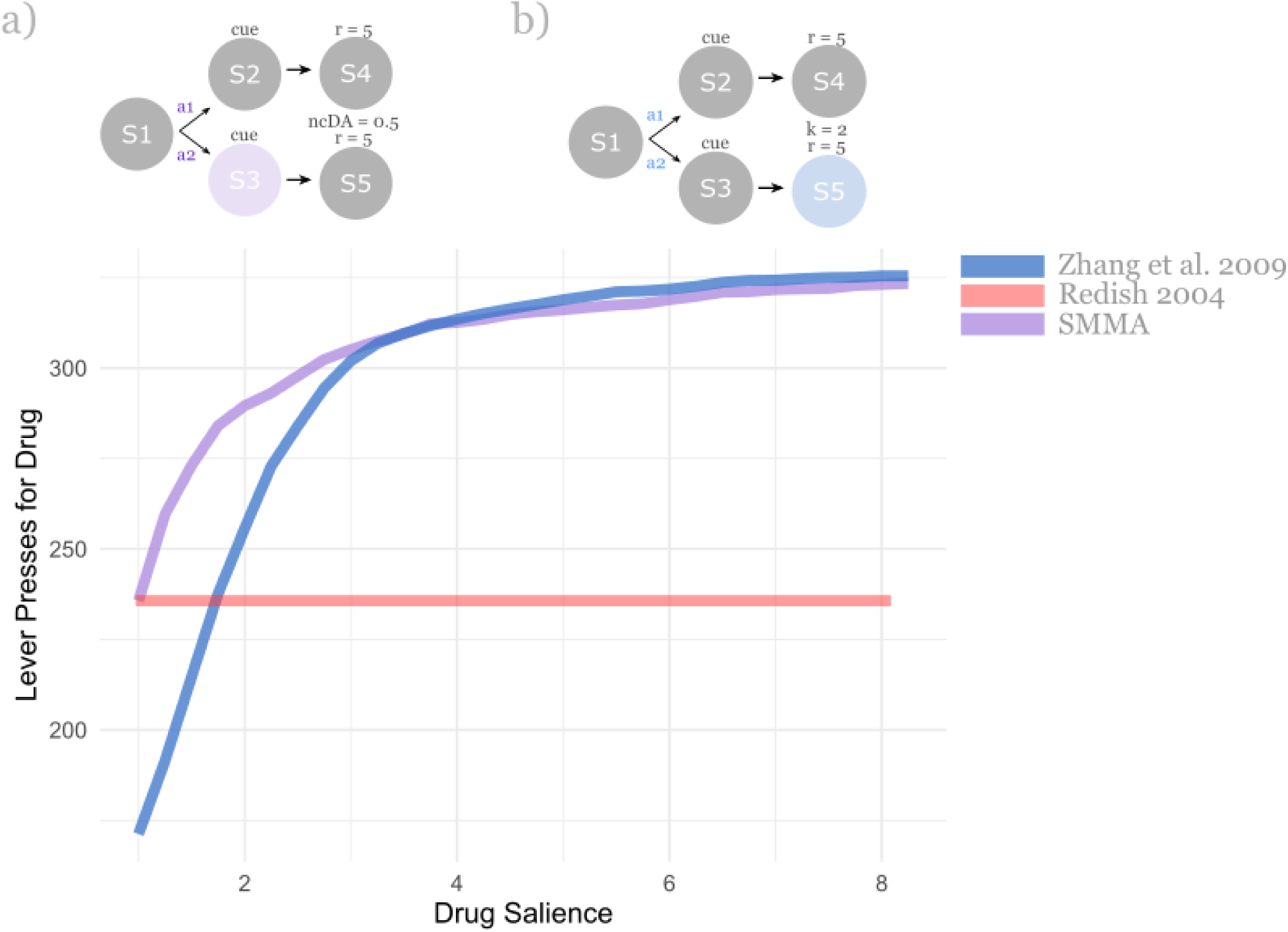
The effect of salience on lever presses for drug rewards. Agents choose between action a1 which led to a non-drug reward, delivered in state 4 (S4) or action a2 which led to a drug reward, delivered in S5. ***a)*** the state space used for the SMMA model simulation, where the salience is applied at the cue state (S3), prior to the drug reward (S5) which is r = 5 and the ncDA = 0.5. The non-drug reward is on S4, where r = 5 and ncDA = 0. ***b)*** the state space used for the Zhang et al. 2009 model, salience (k = 2) is applied to the reward state (S5), representing the drug reward and k = 1 for non-drug reward. The Redish 2004 model does not have a salience factor, and therefore, has no change in behaviour with increasing salience. Both the Zhang et al. 2009 and the SMMA models produced increase in lever presses for the drug rewards (increase in action a2), as the salience was increased.

However, the Zhang et al. 2009 and the SMMA models produce this salience-related effect in different ways. The Zhang et al. 2009 model produces this through salience being multiplied by the reward, so the agent acts as if the reward itself is increased and therefore presses the lever more as salience is increased. The SMMA model produces this result through salience being multiplied by the constant positive prediction-errors due to the drug reward (the ncDA factor). Therefore, as salience is increased in the SMMA model, the weight placed on positive prediction-error is greater and that causes a greater value update and increases in lever presses for the drug reward. A critical conceptual difference between the two models is that the SMMA model predicts that increases in drug behaviours are due to the salience modulating the prediction-errors, but the Zhang et al. 2009 predicts that this is instead due to the salience increasing the magnitude of drug rewards.

### Modelling probability of taking drug actions given contrasting non-drug rewards

A key component of the Redish 2004 model is that the more drug actions the agent takes, the greater the contrasting non-drug reward needs to be in order to reduce the number of drug actions (i.e., the later the stage of addiction, the more difficult it is to choose to abstain through contrasting non-drug rewards). This arises in the Redish 2004 model (Figure 3b) because the value increase is unbounded due to the non-compensable dopamine. This effect does not arise in the Zhang et al. 2009 model because the value plateaus and the prediction-error reaches zero (Figure 3d). However, the SMMA model does produce this effect, due to the inclusion of the non-compensable dopamine component (Figure 3f). In the SMMA model, however, the salience factor interacts with the non-compensable dopamine signal and produces a sharper sigmoid shape (larger beta), and a larger mean of the sigmoid, indicating that a higher contrasting non-drug reward is required to decrease the probability of taking the drug action. Importantly, this interaction between the salience and the ncDA factor accelerates the transition towards the later stages of addiction, where an increasingly greater non-drug reward is required to reduce the probability of taking the drug action. The SMMA model therefore predicts that the progression towards addictive behaviours may be accelerated through this interaction between the high salience for drug cues and the drug dopamine-related positive prediction-errors.

**Figure 3.**
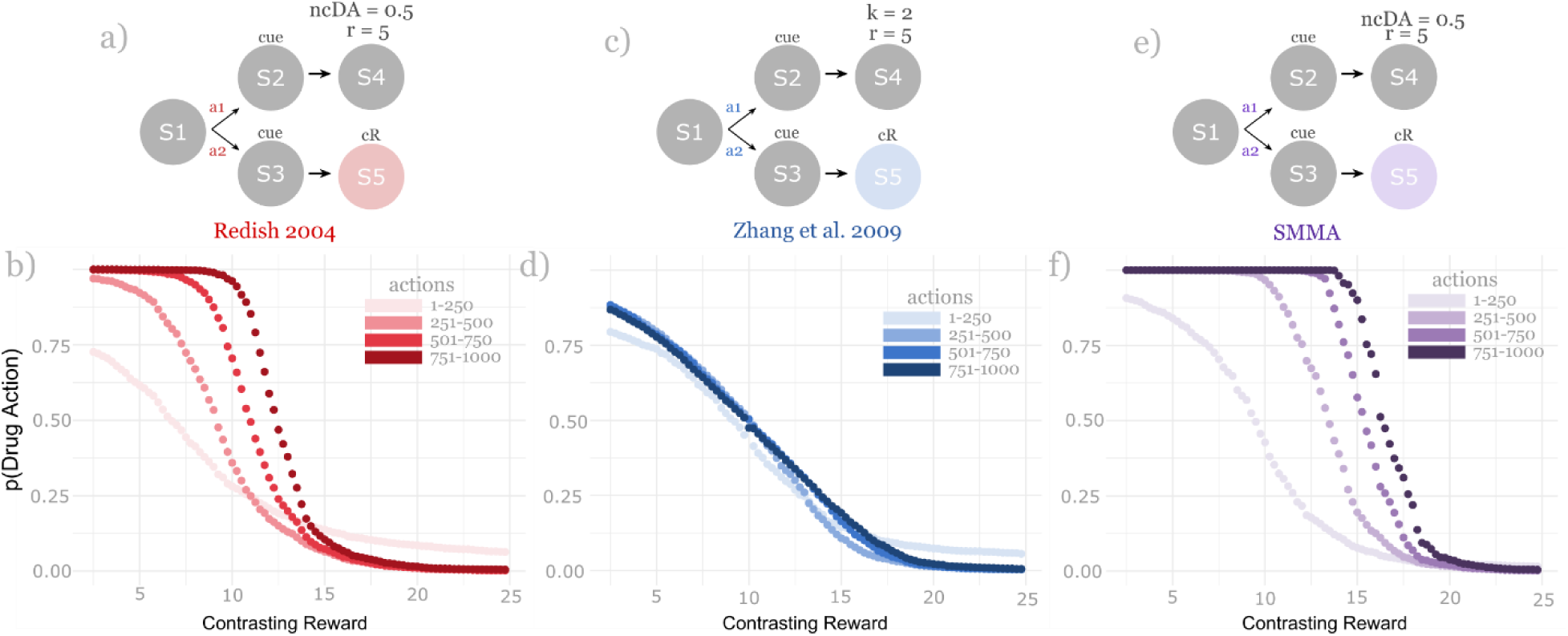
Modelling probability of taking drug actions given contrasting non-drug rewards. In all models, action a2 lead to a non-drug reward, delivered at state 5 (S5) and action a1 lead to the drug reward, delivered in S4. ***a)*** The state space used for the Redish 2004 model, where drug reward is represented as r = 5 and ncDA = 0.5, and non-drug reward has ncDA = 0. ***b)*** as the contrasting non-drug reward was increased, the probability of drug actions (action a1) also decreased. As the number of drug actions taken increases, the contrasting non-drug reward also needs to be increased to reduce the probability of taking the drug action. ***c)*** The state space used for the Zhang et al. 2009 model, where drug reward is represented as r = 5 and salience (k) = 2, and non-drug reward with salience = 1. ***d)*** as the non-drug contrasting reward is increased, the probability of drug actions decreases, however, the number of drug actions taken do not influence the amount of contrasting non-drug rewards needed to reduce the probability of drug actions. ***e)*** The state space used for the SMMA model, where drug reward is represented as r = 5 and ncDA = 0.5, and non-drug reward has ncDA = 0. ***f)*** similar to the Redish 2004 model, as the contrasting non-drug reward increases, the probability of drug actions (action a1) decreases under the SMMA model. Additionally, ss the number of drug actions taken increases; the contrasting non-drug reward needed to reduce the probability of taking the drug action increases. In the SMMA model, the salience and the ncDA factors interact to accelerate this effect.

### Developing inelasticity over time

One of the hallmarks of drug addiction is that people become inelastic to costs over time (Madden & Bickel, 2010). Importantly, while they become more willing to pay high costs for drugs, people with substance use disorders do remain sensitive to those costs, choosing the lower cost options when available (Carroll, 1993). This increased inelasticity to costs over time is captured in the Redish (2004) model due to the unbounded value increase from the ncDA factor (Figure 4b). Increased inelasticity over time is not captured in the Zhang et al. (2009) model due to the value plateauing and prediction-error reaching zero (Figure 4d). However, the SMMA model does produce this developing inelasticity effect over time (Figure 4f). Interestingly, the two effects of salience and the ncDA factor, synergize in the SMMA model to create an even less steep elasticity, because the salience factor accelerates the increase in value for drug rewards, requiring a larger cost to reduce drug actions. The SMMA model therefore predicts that a greater salience towards drug cues could interact with drug dopamine to accelerate the development of inelasticity towards drug-related costs, and as a result there is a persistence in drug behaviours, even when presented with large costs.

**Figure 4.**
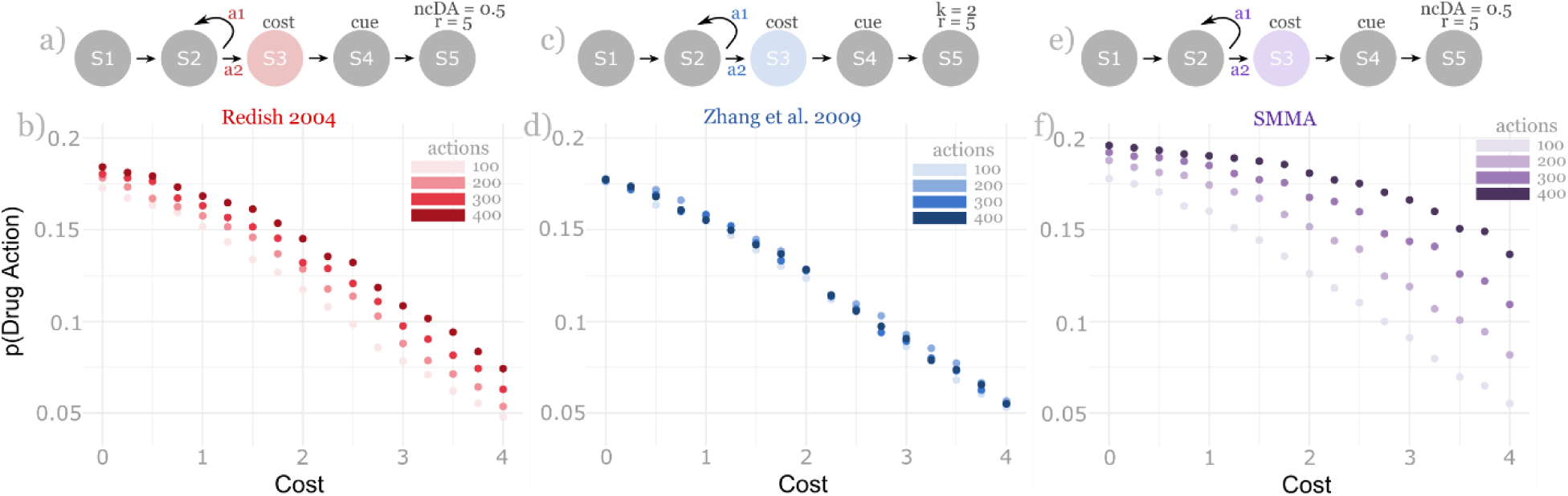
Simulating the development of inelasticity to costs overtime. In all model simulations, the state space was such that action a2 would lead to a given cost in state 3 (S3) but also the drug reward is S5. Action a1 would avoid this cost but would also forgo the drug reward. ***a)*** The state space for the Redish 2004 model, drug reward is given in S5 where r = 5 and ncDA = 0.5. ***b)*** the Redish 2004 shows that agents are sensitive to costs, as greater costs reduces the probability of taking the drug actions, and overtime, as the agents takes more drug actions, they become less sensitive to these costs (develop inelasticity). ***c)*** the state space using the Zhang et al. 2009 model where drug reward is given in S5 where r = 5 and salience (k) = 2. ***d)*** the Zhang et al. 2009 model simulates that as the costs increases, the probability of drug actions reduces, however this is not influenced by the number of drug actions taken, and the development of inelasticity overtime effect is not produced using this model. ***e)*** the state space model for the SMMA model, where drug reward is given in S5, where r = 5 and ncDA = 0.5. ***f)*** The SMMA model also shows that agents are sensitive to costs and become increasingly less sensitive as the number of drug actions they take increases. This is similar to the Redish 2004 model, however in the SMMA model the ncDA and the salience factors interact to accelerate this inelasticity effect.

### Salience attribution on negative and positive prediction-errors using the current SMMA model

People with substance use disorders often continue using drugs despite extreme negative outcomes. This is often a large component in defining substance use dependence (*Diagnostic and Statistical Manual of Mental Disorders (5th Ed.)*, 2013). In the SMMA model, negative prediction errors have a reduced salience prior to reaching drug states, especially when compared to before reaching non-drug states (Figure 5g). The opposite is true for positive prediction-errors, which are weighted with more salience compared to the positive prediction-errors prior to reaching non-drug reward states (Figure 5i). Therefore, if the agent is in a high dopamine state, the cost that occurs before reaching the drug state is downweighted, and the reward is overweighted. These effects makes the agent more likely to take actions leading to the drug state. This asymmetric learning effect cannot be produced using the Zhang et al. (2009) model as negative and positive rewards are weighted equally in salience (Figure 5c and 5e). It also cannot be produced using the Redish (2004) model as there is no salience factor and the model cannot produce a negative prediction-error, for either drug and non-drug rewards (Figure 5b and 5d).

**Figure 5.**
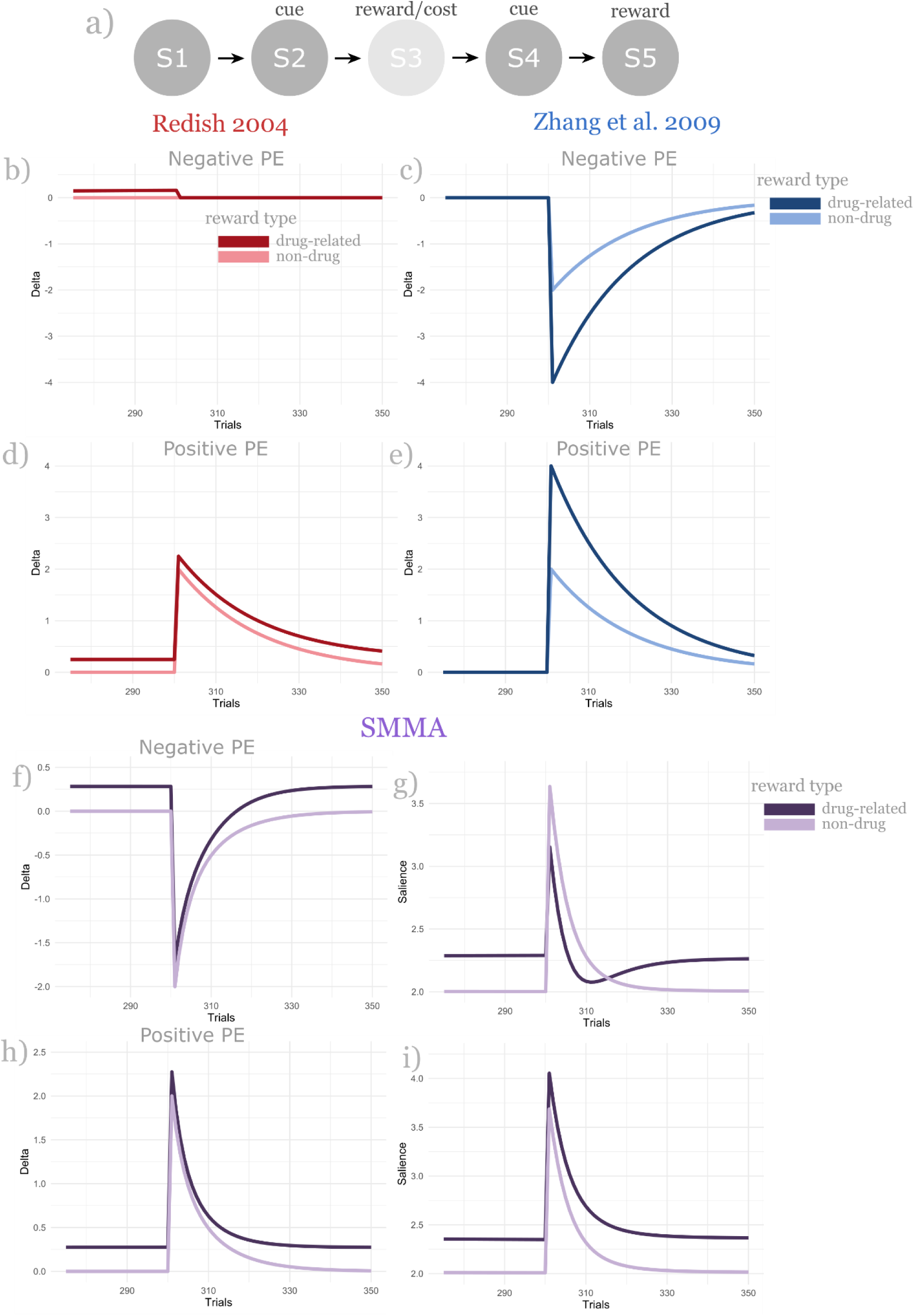
Weighting positive and negative reward prediction-errors from drug-related and non-drug rewards. ***a)*** the state space used for all three models used for the simulation. The agent gets a cue in state 2 (S2) and then a non-drug reward or cost in S3, which is where the positive or negative prediction-error is generated, and all plots are from this state. S4 is another cue state, and in state 5 agents either receive the drug or the non-drug reward. ***b)*** the drug-related and non-drug negative prediction-error using the Redish 2004 model. Because the minimum prediction-error allowed under the Redish 2004 model is 0, there is no negative prediction-error produced here for both drug-related and non-drug related rewards. ***c)*** the drug-related and non-drug negative prediction-error under the Zhang et al. 2009 model. Due to the drug-related reward having a greater salience, negative prediction-errors here are more negative for drug-related rewards than for non-drug rewards. ***d)*** and ***e)*** the positive prediction-errors from both drug-related reward and non-drug reward using the Redish 2004 and the Zhang et al. 2009 model, respectively. Both models have higher positive prediction-errors for drug-related rewards than for the non-drug reward. ***f)*** drug-related and non-drug negative prediction-errors using the SMMA model. Non-drug negative prediction-error is more negative for than for drug-related reward. ***g)*** the salience placed on the drug-related negative prediction-error is lower than the salience placed on the non-drug negative prediction-error. ***h)*** drug-related and non-drug positive prediction-errors using the SMMA model. Drug-related positive prediction-error is more positive than non-drug. ***i)*** the salience placed on drug-related positive prediction-errors are greater than the salience placed on the non-drug positive prediction-errors.

In the SMMA model, this asymmetric internal representation updating effect arises from the average prediction-error being greater for drug-related rewards than for non-drug related rewards. An increased average prediction-error is driven by the continual positive prediction-errors caused by the ncDA factor. With a greater average prediction-error, the salience on the negative prediction-error is downweighted and overweighted on the positive prediction-error.

Another factor influencing salience weights is that the negative prediction-error is slightly lower for the drug-related negative prediction-error (Figure 5f) but greater for the positive prediction-error (Figure 5h). This is also caused by the ncDA factor, where the drug states have a constant positive prediction-error and so any prediction-error is using this positive-prediction error as the starting point and causes the negative prediction-error to be less negative and positive prediction-errors more positive for drug rewards because it starts at a prediction-error already above 0. Overall, the positive ncDA for drugs has two consequences; 1) it produces a greater average prediction-error for drug-states, and this causes the different weights placed on the salience of negative and positive prediction-errors under our salience formula and 2) it produces a positive baseline prediction-error for drug states, causing different magnitudes in the prediction-errors. Together, these factors explain a reduced salience and updating from any negative prediction-errors in drug states but at the same time, an increased salience on any positive prediction-errors in drug related states. Importantly, this produces a misaligned internal representation where the agent learns/updates more from positives, but downweights the negatives prior to receiving the drug reward.

Going further, we simulated whether the average prediction-error factor alone could influence the salience placed on positive and negative prediction-errors, without any influence of ncDA. Figure 6 shows the salience placed on a prediction error of +5 and −5. As modelled using our mathematical definition of salience (equation 8), positive prediction-error has more salience as the average prediction-error increases. And the opposite is true for the negative prediction-error, which has less salience, as the average prediction-error increases. Therefore, the SMMA model, with its mathematical salience definition, is not dependent on the ncDA factor alone to produce these misaligned internal representations, but any factor that increases the average prediction-errors. The ncDA factor is one possible way in which drugs can increase the average prediction-error and cause this salience misattribution effect, but not the only possible factor.

**Figure 6.**
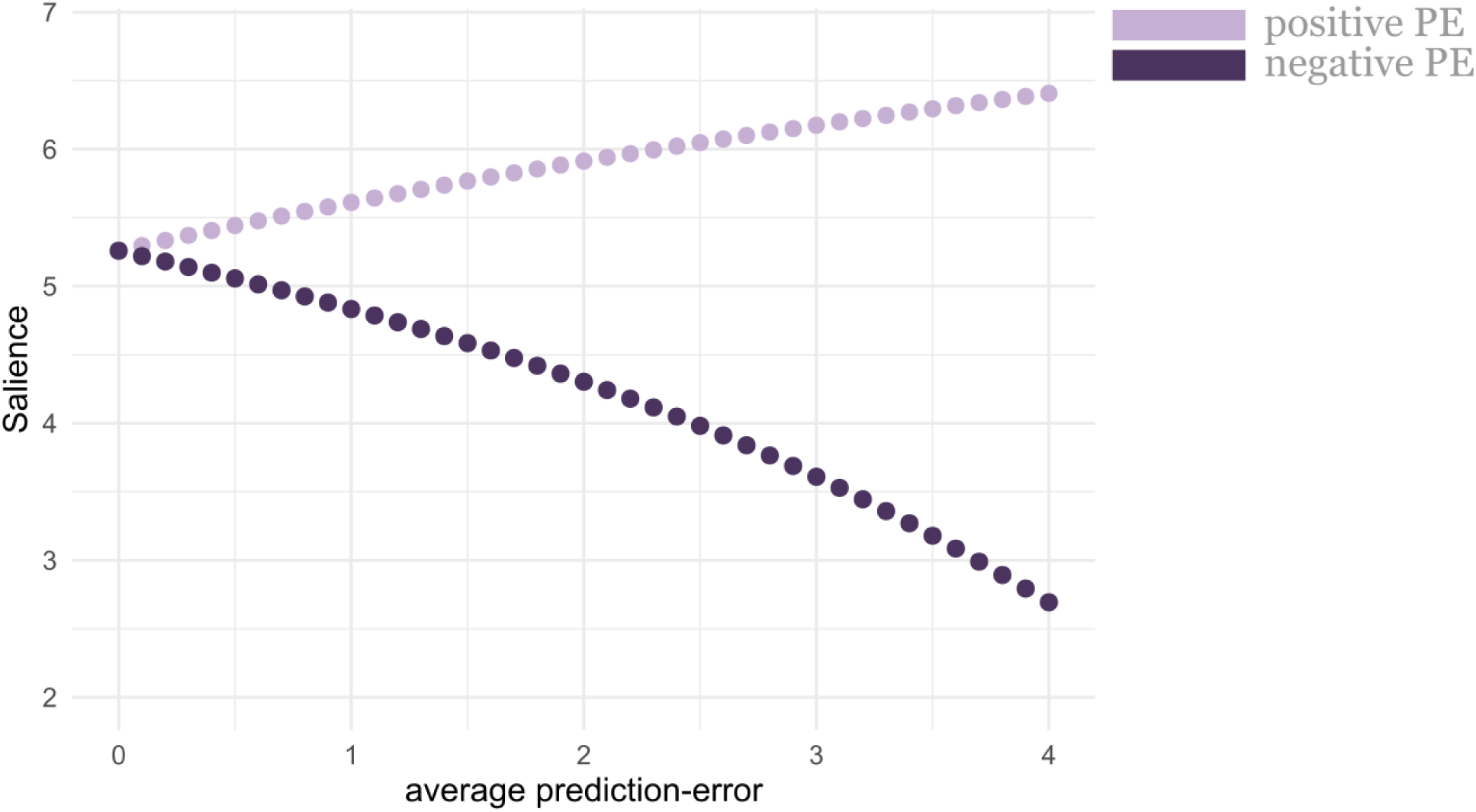
The effect of increasing average prediction-error on the salience placed on positive and negative prediction-errors. The salience placed on negative and positive prediction-errors as the average prediction-error increases. The greater the average prediction-error, the more salience on the positive, and less on negative. Therefore, as an agent is in an increasingly high dopamine state (high positive prediction-error rate), positives are used more to update the internal representations and the negatives are used less to update the internal representation. PE = prediction-error.

### Reversal learning with drug and non-drug rewards

Aberrations in reversal learning tasks are generally found in people with substance use disorder (Verdejo-Garcia et al., 2018). Here, we tested all three models in a reversal learning simulation, with the non-drug cost/reward given prior to the drug reward or another non-drug reward. The state-space model (Figure 7a-b) here was designed to simulate a situation where prior to receiving the drug-reward, the agent would either get a cost or a reward, which is reversed after several trials, but the drug reward itself would not be reversed (i.e., the model changes the pre-signal for drug-seeking where if you now get punished instead of rewarded for taking the drug actions, do you switch your behaviour towards the non-drug action?). Negative consequences of drug-seeking are generally more prominent in later stages of addiction (Brand et al., 2016, 2019; Robinson & Berridge, 1993), and as a result we started this simulation with drug-seeking being rewarded (e.g., the initial social reward), and then reversed to now being preceded with a cost.

**Figure 7.**
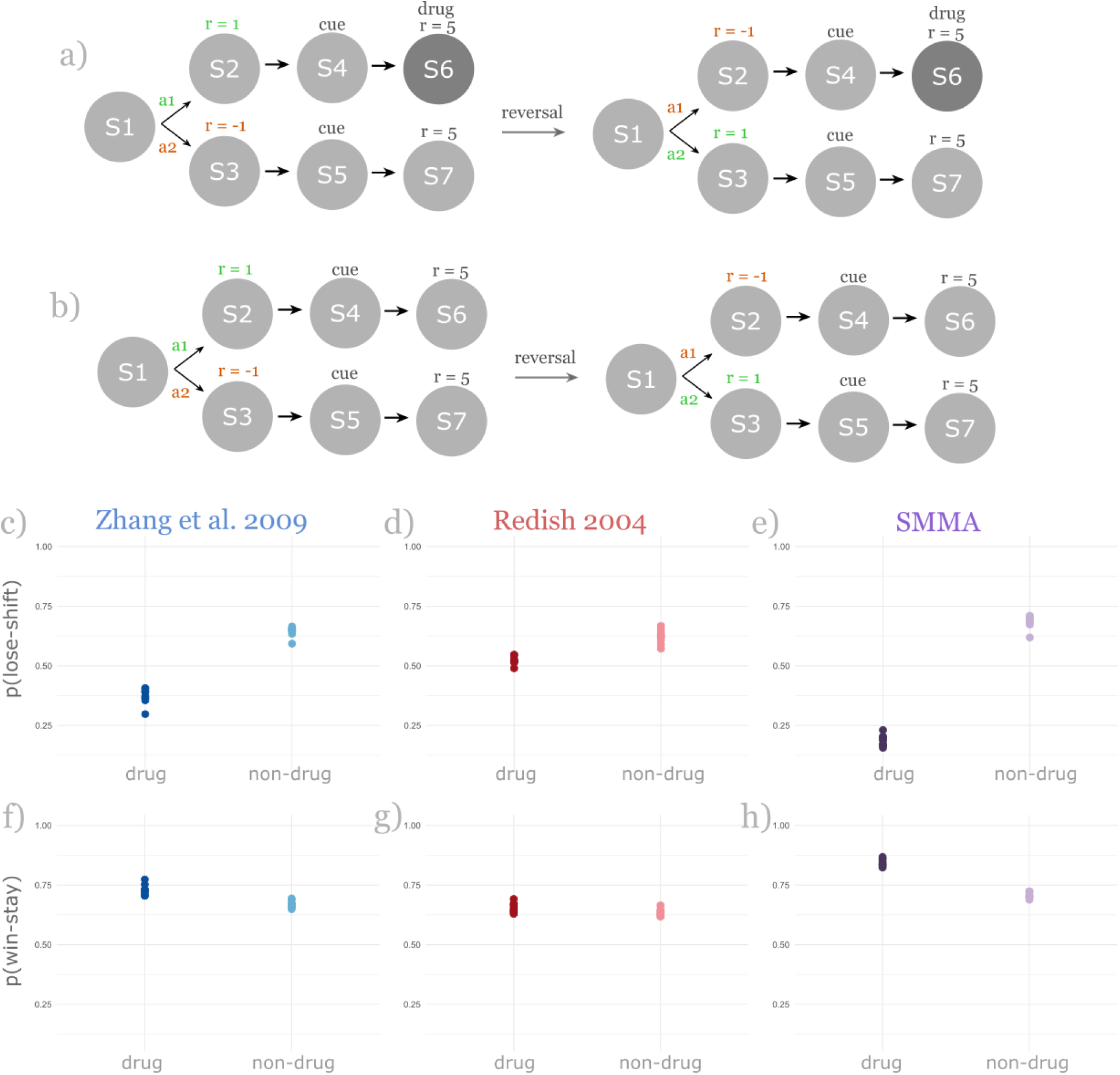
Reversal learning from drug and non-drug related costs and rewards. ***a)*** the state-space model used for the drug-related costs and reward simulation. Here, action a1 leads to a reward (r = 1) in state S2, followed by a drug reward in S6. In the SMMA and the Redish 2004 model, drug reward is represented by r = 5 and ncDA = 0.5. In the Zhang et al. 2009 model, drug is represented as salience (k) = 5, and non-drug with salience = 1. If the agent chooses action s2, it will receive a cost in S3 (r = −1), followed by a non-drug reward in S7. When there is a reversal (after every 100 trials), action a1 now leads to a cost in S2, but the agent will still receive the drug-reward in S6. ***b)*** the state space used for the non-drug simulation. Here if the agent took action a1, it would get a reward (r = 1) in S2 and another reward (r = 5) in S6. If the agent chose action a2, it would get a cost (r = −1) in S3 and a reward (r = 5) in S7. There is a reversal after every 100 trials, where the cost is now in S2 and reward in S3, all else remains the same. In both, drug and non-drug simulations, taking the action that leads to the cost is defined as a loss, and win as the state opposite to this. ***c)*** the probability of lose shift (p(lose-shift)) using the Zhang et al. 2009 model. Agents had a lower mean lose-shift probability for the drug simulation (0.37) compared to the non-drug simulation (0.64). ***d)*** the Redish 2004 followed the same trend with mean p(lose-shift) for the drug simulation (0.52) being less than the non-drug simulation (0.62). ***e)*** the SMMA model also had a lower mean p(lose-shift) for the drug simulation (0.19) than the non-drug simulation (0.69). ***f)*** the probability of win stay (p(win-stay)) using the Zhang et al. 2009 model. Agents had a high mean win-stay probability for the drug simulation (0.73) compared to the non-drug simulation (0.67). ***g)*** the Redish 2004 followed the same trend with mean p(win-stay) for the drug simulation (0.65) being more than the non-drug simulation (0.63). ***h)*** the SMMA model also had a higher mean p(win-stay) for the drug simulation (0.84) than the non-drug simulation (0.71). The inverse temperature parameter used here was 0.25 instead of 0.5, this to avoid ceiling effects.

All three models demonstrated a higher probability of getting a reward and repeating this rewarding action (win-stay) if that reward was followed by a drug reward, compared to if it was followed by another non-drug reward (Figures 7f-h). This is due to the drug states having a higher value in all three models, compared to the non-drug states. The SMMA model has the highest win-stay probability for drug states, compared to the other models for two reasons, 1) the reward prior to the drug reward is overweighted in salience and therefore has a greater positive prediction-error and value update, and 2) the ncDA and the salience factors interact to accelerate the value increase of the drug state. In the Zhang et al. 2009 model, only the drug reward is increased in salience, and the reward preceding this is not changed in magnitude. The Redish 2004 model also treats the reward prior to the drug reward to be unchanged in magnitude, even as the drug state increases in value, due to the ncDA factor. Therefore, a critical difference between the models is that the SMMA model overweights the reward for drug-seeking in addition to increasing the value for the drug state, whereas the Redish 2004 and the Zhang et al. 2009 models only overweight the value of the drug state.

All three models also demonstrated that the agent becomes less likely to shift strategy after receiving a cost (lose-shift) if the drug reward is received after this cost, compared to if a non-drug reward is received after the cost. Therefore, in all three models, agent appear less sensitive to costs if that cost is followed by a drug reward, due to this lower lose-shift probability (Figure 7c-e). However, we see this effect in the Redish 2004 model and the Zhang et al. 2009 model primarily due to the value of the drug state being much greater than the value of the non-drug state, and not because the cost prior to the drug states is downweighted in magnitude. In the SMMA model, however, this effect occurs for two reasons: 1) the negative prediction-error generated due to the cost prior to the drug reward is downweighted in salience and therefore the value of the drug-seeking state is less negative, and 2) the drug state has a greater value due to the ncDA and the salience factor interacting to accelerate the value increase. Therefore, the ncDA and the salience factors in the SMMA model synergise and produce a very low lose-shift probability for drug actions, by both, reducing the salience placed on the cost for drug-seeking and by accelerating the increase in the value update of the drug state itself. In sum, the SMMA model predicts that the costs for drug-seeking is downweighted, and the value of the drug state is overweighted. This is in sharp contrast to the Zhang et al. 2009 model and Redish 2004 model, which do not downweight any costs for drug-seeking, but only overweight the value for the drug state.

### Reversal learning with non-drug rewards and variable average prediction-errors; the link with behavioral addictions

Behavioural or non-pharmacological addictions such as gambling, social media, and video/online gaming show similar behavioural deficits to drug addictions (Brand et al., 2019; Chamberlain et al., 2016; Ognibene et al., 2019; L. Wei et al., 2017). In the SMMA model, a change in average prediction-error can produce reduced sensitivity to learning from negative prediction-errors and increased sensitivity to learning from positive prediction-errors without any influence from the ncDA factor (see Figure 6). We tested how the asymmetric learning effects could produce problematic behaviours, even without a ncDA drug factor.

In the reversal learning simulation here, win-stay captures sensitivity to positive prediction-errors and lose-shift captures sensitivity to negative prediction-errors. In Figure 8b, as the average prediction-error is increasing, there is initially a dip in the likelihood of staying after a win. This dip is a consequence of using the log scale in the salience formula, where the salience on the negative prediction-error decreases faster than the salience on the positive prediction-error increases (see Figure 8d). After the average prediction-error increases above 2, there is a continual increase in the probability of win-stay with increasing average prediction-error as the net salience and value update of the win state is high enough and the value of the lose state is low enough for the agent to win and stay in the win state.

**Figure 8.**
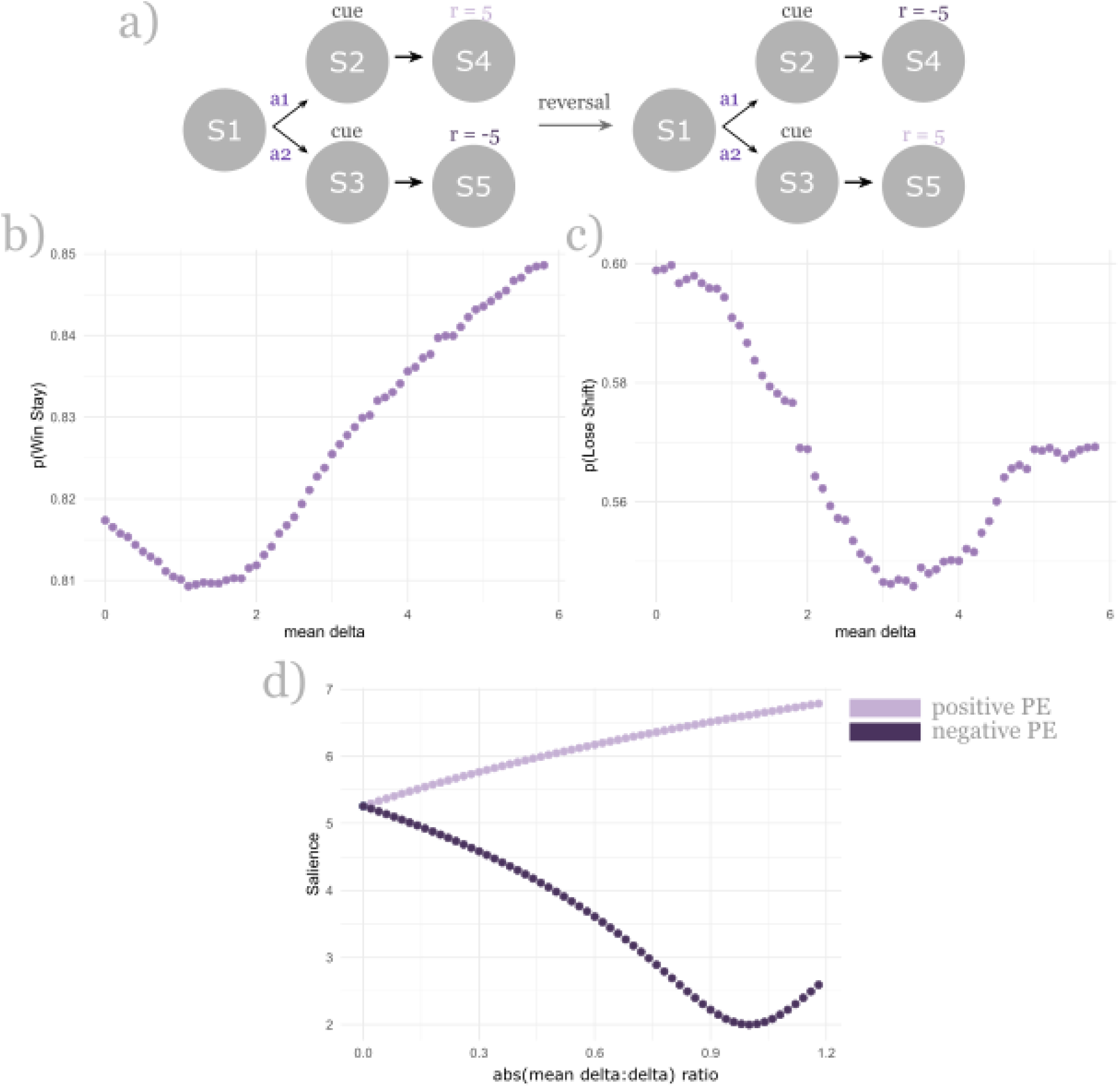
The influence of average prediction-errors on reversal learning, without the ncDA factor. ***a)*** the state space used, where there is a reversal in cost/reward in S4 and S5. ***b)*** as the average prediction-error increases, the probability of win stay does too, after the initial decline caused by using the log scale where positives increase less than the negatives decrease. ***c)*** The probability of lose-shift decreases as the average prediction-error increases, with an upper bound, in which case it increases. ***d)*** the salience placed on the positive prediction-error increases as the average prediction-error increases, and the salience on negative prediction-error decreases, but at a faster rate than the positive increases (due to the log scale). After the absolute ratio of the average prediction-error: prediction-error reaches 1, the upper bound limit is reached and the salience on negative prediction-error increases. The salience on the positive prediction-error increases without a bound. *Abs = absolute*.

Next, the simulation demonstrated a decrease in the probability of leaving after a loss as the average prediction-error increased (Figure 8c). This effect arises from the reduced salience on the negative prediction-errors as the average prediction-error increases – causing the value on the lose state to be less negative and therefore making the agent less likely to change strategy (lose-shift). However, this drop only lasts as long as the negative prediction error is equal to or less than the average prediction-error (i.e., when the absolute ratio of average prediction-error and the negative prediction error is 1 or less). This is because in the current salience formula, the salience is lowest when the prediction-error plus the average prediction error is close to 0 (e.g., a prediction-error of −2.5 will have the lowest salience when the average prediction-error is 2.5, where the absolute ratio is equal to 1 and the sum of the two is 0). When the absolute ratio becomes greater than one (i.e., where 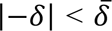), the salience on the negative prediction-error starts to increase

(Figure 8d), and so does the probability of leaving after a loss – as now the lose state becomes increasingly negative in the value, making shifting more favourable. The greater the average prediction-error, the less reliable the internal representation (i.e., the model is making a lot of inaccurate predictions and is therefore getting a large number of prediction-errors). When the average prediction-error is so large relative to the instantaneous negative prediction-error, to the point that the absolute ratio between the two is greater than one 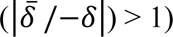, the salience is no longer downweighted on these negative prediction-errors. Therefore, the effect of downweighting salience on the negative prediction-error is bounded by 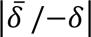 being less than 1.

In sum, as the average prediction-error increases, the model shows an increase in win-stay behaviours, indicating a greater sensitivity to reinforcement (albeit after the initial decline due to the log scale). There is also a decrease in lose-shift behaviours as the average prediction-error increases, indicating a reduced sensitivity to aversion. However, this decrease in lose-shift behaviours is bounded by the absolute ratio of 1 or less between the average prediction-error and the negative prediction-error. This asymmetric learning effects arises from the mathematical definition of salience under the current SMMA model; neither the Zhang et al. 2009 nor the Redish (2004) model produce this effect (see Figure 5). These results overall suggest that a high average prediction-error rate may be one possible predictor or contributor to the development of behavioural addictions, where there is a strong persistence in the behaviour despite negative consequences.

### Delay discounting simulations

People with a substance use disorder generally discount drug rewards faster than non-drug rewards (Bickel & Marsch, 2001). Neither the Zhang et al. 2009, nor the Redish 2004 model can adequately produce this effect, with only a very slight steeper discounting effect for drug rewards in the Redish 2004 model, and none at all with the Zhang et al. 2009 model (Figure 9b and 9d). However, the SMMA model did produce this effect where drug rewards were discounted much faster than non-drug rewards (Figure 9f). The mathematically defined salience factor form the SMMA model was necessary to produce steeper discounting for drug rewards, the ncDA factor alone cannot produce this.

**Figure 9.**
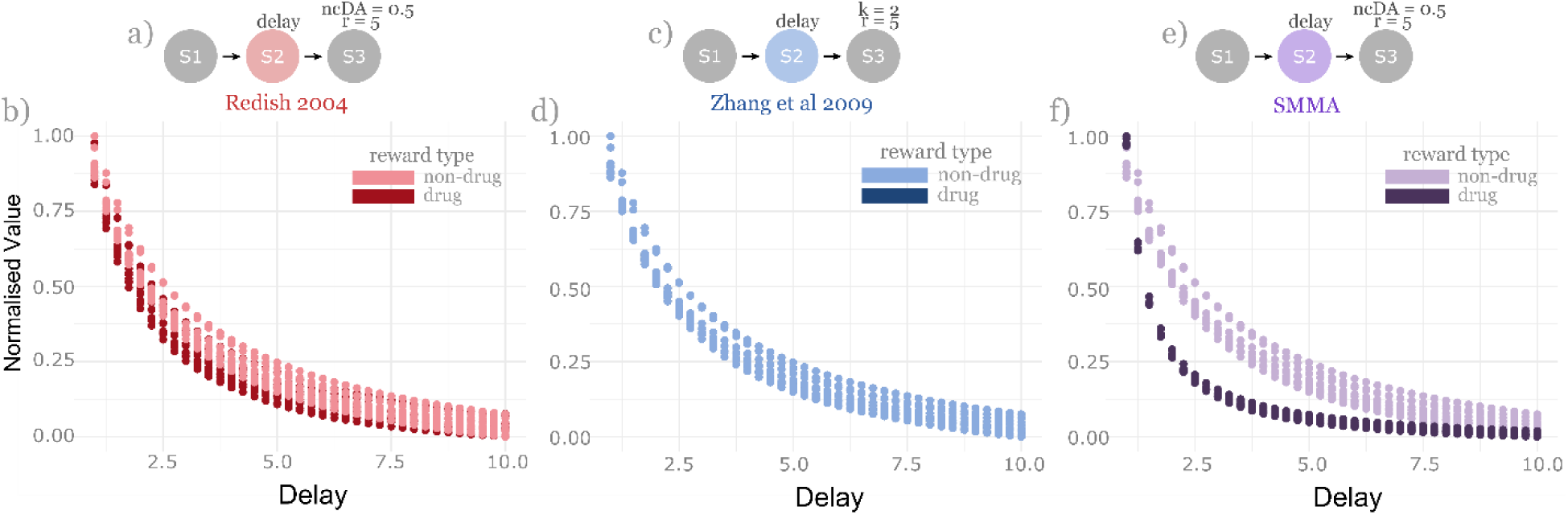
Delay discounting curve for drug and non-drug rewards using all 3 models. In all simulations, agents had a variable delay in state 2 (S2), with the drug or non-drug reward delivered in state 3 (S3). ***a)*** the state space used for the Redish 2004 model where drug reward was r = 5 and ncDA = 0.5 and non-drug reward was r = 5. ***b)*** the drug rewards are only very slightly discounted steeper than non-drug rewards under the Redish 2004 model. ***c)*** the state space used for the Zhang et al. 2009 model where drug reward was r = 5 and salience (k) = 2, and non-drug reward was r = 5. ***d)*** the drug and non-drug rewards are discounted identically under the Zhang et al. 2009 model. ***e)*** the state space used for the SMMA model where drug reward was r = 5 and ncDA = 0.5 and non-drug reward was r = 5. ***f)*** the drug rewards are discounted much faster compared to non-drug rewards under the SMMA model.

High dopamine levels contribute to steeper delay discounting (Ballard et al., 2015; Pine et al., 2010). We simulated delay discounting with a variable average prediction-error for drug rewards, representing different dopamine levels, to further investigate how the ncDA and the salience factor interact to produce discounting behaviours. As the average prediction-error increased (higher dopamine states), the model showed steeper discounting for drug rewards (Figure 10), however, this effect was not produced for non-drug rewards (without the ncDA factor). Therefore, the two simulations suggest that, both the ncDA and the salience factors are necessary to produce the steeper delay discounting effect. The ncDA factor causes an unbounded value increase for drug rewards, which the salience factor then modulates by placing less salience on the delay as the average prediction-error increases, and thereby reducing the value of drug rewards as time to the drug reward increases. The SMMA model thereby predicts that the reduced salience placed on time, as the agent is in a higher dopamine state, is one contributor to the steeper discounting of drug rewards found in people with a substance use dependence.

**Figure 10.**
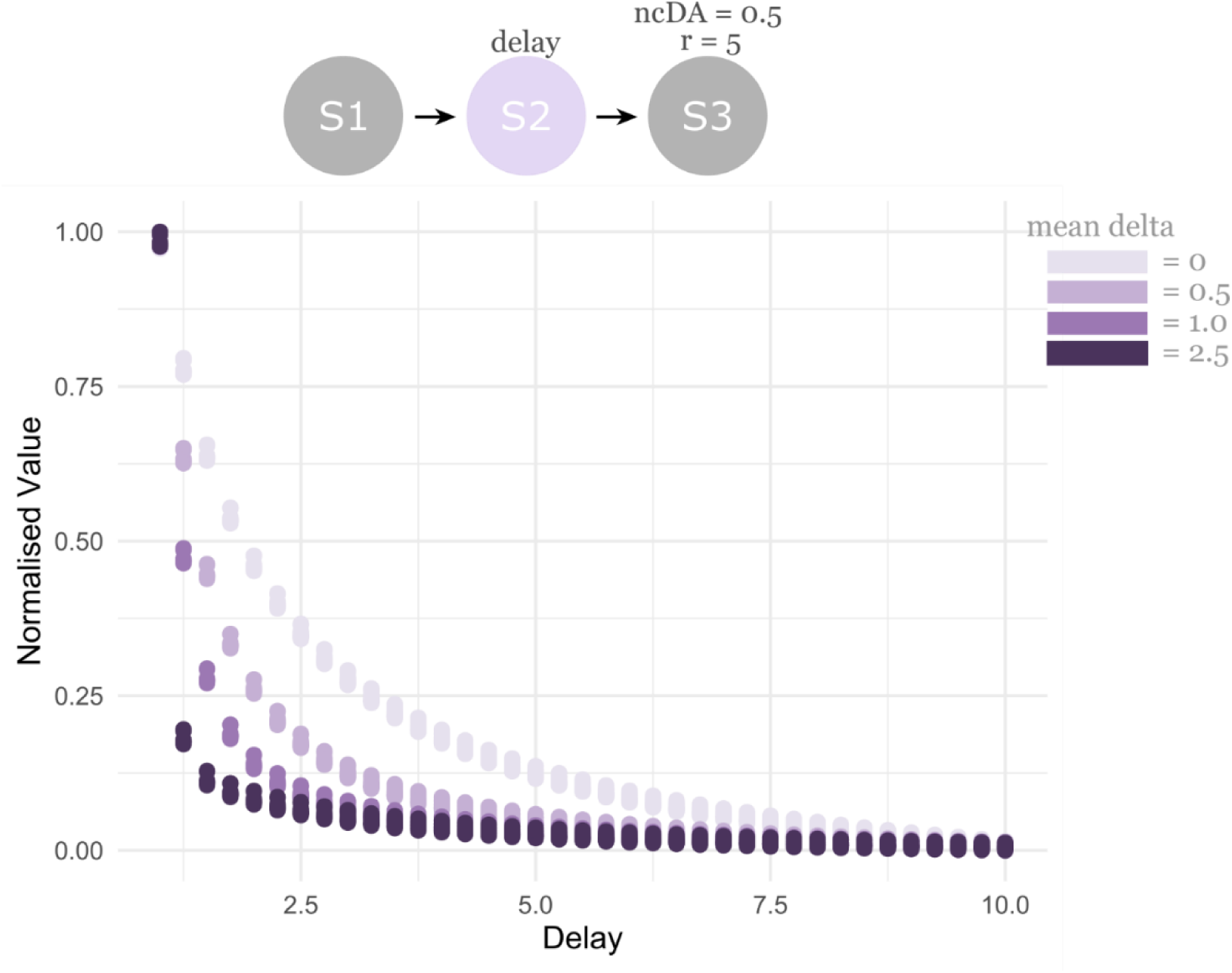
The effect of average prediction-error on delay discounting of drug rewards using the SMMA model. As the average prediction-error increases for drug rewards, there is a steeper discounting of the drug reward.

### Induced craving, bingeing, and salience simulations

Craving influences value multiplicatively, and not additively or exponentially (Konova et al., 2018). Here, we operationalised craving as a state where the average prediction-error is negative. This means that it can influence the salience placed on prediction-errors, and as a result it also influences valuation multiplicatively under our model, using equation 9 to update values, consistent with the data from Konova et al., (2018). We induced a craving state in our agents through two steps. The first step involved generating the expectation of receiving the drug reward (by allowing the agent to get the drug for 30 trials). Following this, the drug reward was no longer given, causing negative prediction-errors to the point where the mean prediction-error became negative (by trial 75; 45 trials after drugs were removed). At this point (when mean prediction-error was negative), either a non-drug negative or a non-drug positive prediction-error (of 1.75) was given, and the salience placed on these negative and positive-prediction errors were plotted. We are therefore simulating how an agent weights non-drug positive and negative prediction-errors, while in a craving state. When the agent is in an induced craving state, where the average prediction-error is negative, it down-weights salience on a non-drug positive prediction-error (Figure 11a) but overweights salience on a non-drug reward negative prediction-error (Figure 11b). Therefore, the model predicts that a craving state may lead to downweighting any positives, but overweighting any negatives, which is a component of depressive-like symptoms (Rouhani & Niv, 2019).

**Figure 11.**
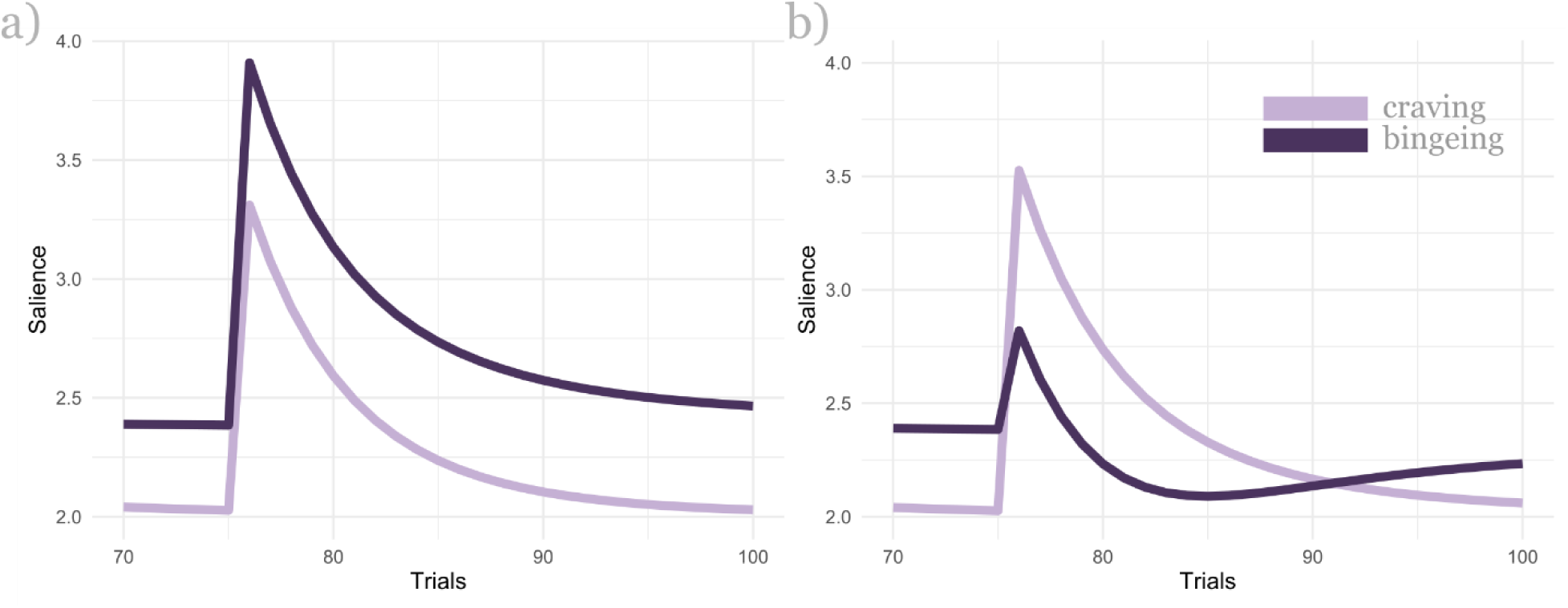
The salience placed on negative and positive prediction-errors under induced craving and bingeing states. Under the induced caving state, ***a)*** the agent places less salience on positive prediction-errors and ***b)*** more salience on negative prediction-errors.

People with a substance use disorder also show steeper delay discounting when in a craving state (Ashare & Hawk, 2012; Giordano et al., 2002). To simulate this effect in the SMMA model, craving was operationalised as a state where the current average prediction-error is negative (i.e., the agent is expecting drugs but does not receive it, eliciting negative prediction-errors to the point where the average prediction error is negative). A negative prediction-error state could be achieved through learning, where there are constant negative prediction-errors, as is the case in the above simulation. However, dopamine depletion, developed tolerance or dopamine antagonists also contribute to a low dopamine state (Jackson-Lewis & Przedborski, 2007; Volkow et al., 1997; Volkow & Li, 2004; Woolverton & Virus, 1989) and therefore may also cause a negative prediction-error state, without any re-learning. At this negative average prediction-error craving state, in the model, there is steeper discounting for drug rewards, compared to when the average prediction-error is 0 (Figure 12). Importantly, this only happens for drug rewards (which comes with a positive ncDA), not the non-drug rewards. Therefore, to produce the steeper delay discounting effect here, the model is depended on the interaction between the salience and the ncDA factors. The negative average prediction-error causes a reduced salience to be placed on time, which in turn reduces the unbounded value updates (caused by the ncDA factor) as the delay is increased. Overall, the SMMA model predicts that there is steeper discounting for drug rewards in a craving state due to a negative average prediction-error state, which causes a lower salience to be placed on the delay to the drug reward, and therefore downweights value as the delay to the drug reward increases.

**Figure 12.**
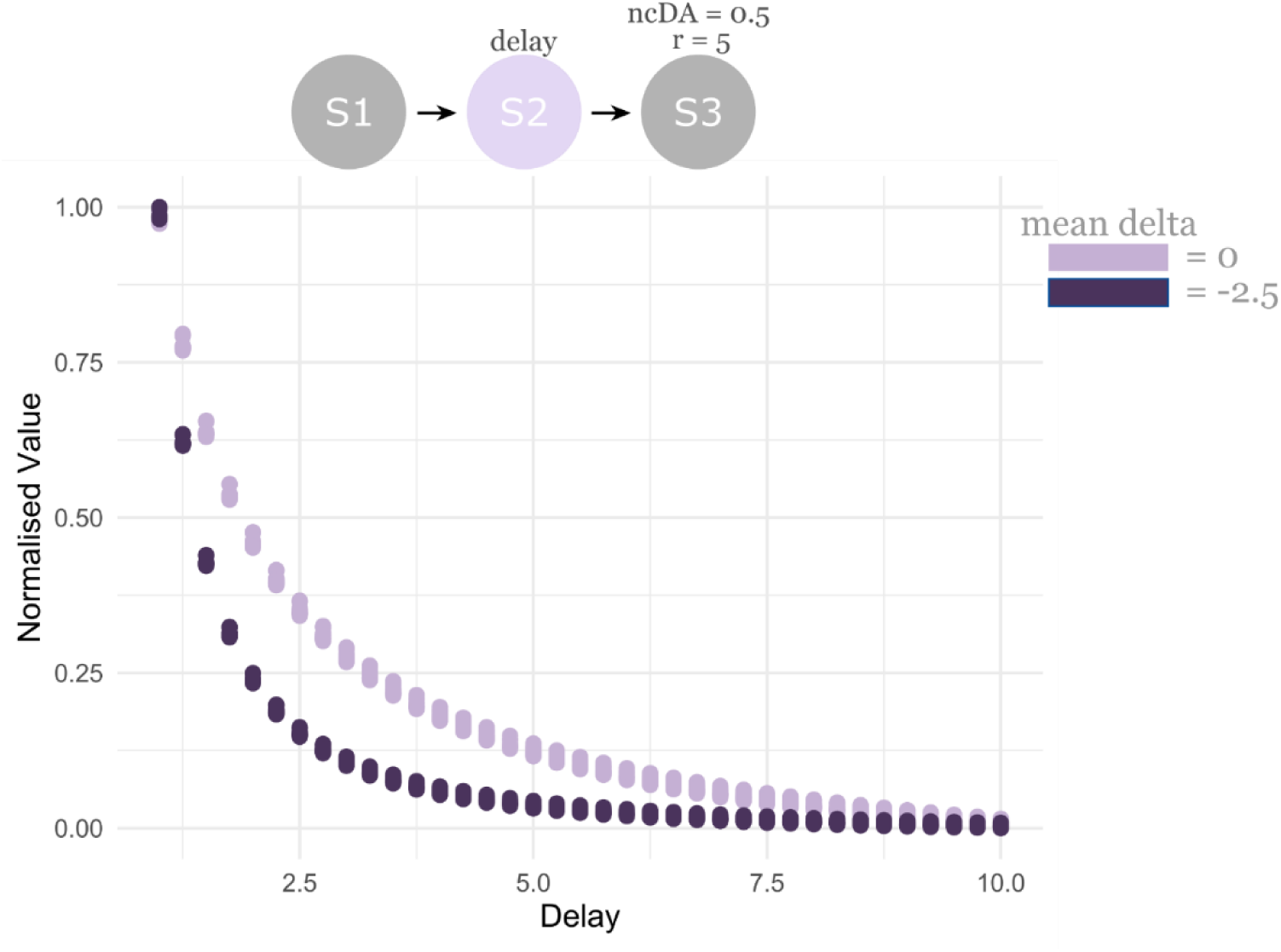
Delay discounting for drug rewards under craving and non-craving states. There is steeper discounting for drug rewards under a craving state, which is when the average prediction-error is a negative, relative to when the average prediction error is 0.

## DISCUSSION

The Redish 2004 model was built on the reward-prediction error theory of dopamine by adding a non-compensable dopamine factor to the delta equation (Redish, 2004). The non-compensable dopamine factor caused constant positive prediction-errors due to drug taking and reinforced addictive behaviours as a result. The Zhang et al. 2009 model was built on the *incentive salience* theory for addiction by adding a salience factor multiplying the reward term in the delta equation (Berridge & Robinson, 2016; Robinson & Berridge, 1993; Zhang et al., 2009b). The greater salience here represented the higher motivational salience value of drugs, and reinforced drug actions as a result. Both models, though conceptually very different, reinforce drug behaviours through value increases of the drug states. Both models present limitations. They do not explain the reduced sensitivity to drug-related costs/negative consequences, the increased impulsivity generally found in people with a substance use disorder, and craving behaviours – all of which are key hallmarks of addictive behaviours. We addressed these limitations by proposing a new model, which we refer to here as a salience misattribution model for addiction (SMMA, see also Kalhan et al., (2021)). The SMMA model is a combination of the Redish 2004 and the Zhang et al. 2009 models, with a novel mathematical definition of salience. Under our model, drug-related negative prediction-errors are downweighted in salience, but at the same time, salience on drug-related positive prediction-errors are overweighted. In SMMA, drug rewards are also discounted faster than non-drug rewards, owing to a lower salience placed on the delay to drug rewards. The SMMA model produces asymmetric learning due to this salience misattribution effect and consequently, a misaligned internal representation is formed. Within this misaligned internal representation, the positives of drugs are misattributed with a greater salience than the negatives, causing reinforcement of drug actions. Using a single parameter regime, we simulated addictive behaviours that the Zhang et al. 2009 and Redish 2004 models also produce but we went further in simulating: 1) downweighting of drug-related negative prediction-errors, 2) steeper delay discounting of drug rewards, 3) craving behaviours and 4) aspects of behavioural/non-pharmacological addictions. This builds on the conceptual framework that salience modulates internal representation updating and may contribute to addictive behaviours by producing misaligned internal representations (Kalhan et al., 2021).

### Does salience-related dopamine modulate learning or motivation?

A key question that arises from the proposed model is whether the salience factor modulates motivational processes where a change in behavior is produced without any relearning (performance), as the Zhang et al. 2009 model suggests. Alternatively, salience may play a role in modulating learning, which is an incremental process, consistent with the Redish 2004 model. The answer is not straightforward, as it likely involves both processes. The SMMA model provides an explanation for cue-triggered fluctuations in behaviors, where a drug cue releases dopamine such that the average prediction error is suddenly increased, and drug behaviors are reinforced through the increased value of drug states. This interpretation is consistent with recent work, where dopamine is viewed as enhancing performance/motivation through increasing the value of the current action (Berke, 2018; Hamid et al., 2015). However, the SMMA model also treats salience as a modulator of learning and internal representation updating, where the lower salience of negative prediction-errors decreases learning, and the higher salience of positive prediction-errors increases learning. The theoretical layout of the SMMA model aligns more closely with the concept of salience modulating learning, by weighting the reward prediction-error learning/updating signals (Keiflin & Janak, 2015; Schultz et al., 1997; Steinberg et al., 2013). This learning interpretation fits well with data suggesting that dopamine modulates neural plasticity by facilitating late long-term potentiation and may increase addictive behaviors through these long term learning and memory mechanisms (Berke & Hyman, 2000; Hyman, 2005; Li et al., 2003; Lisman et al., 2011; Mockett et al., 2004; Sajikumar & Frey, 2004). However, there is a strong possibility that the salience factor in the current model is also playing a role in motivation/performance.

Traditionally, the dissociable roles of dopamine in learning and motivation were thought to be based on the slow ‘tonic’ and fast ‘phasic’ dopaminergic function. The slower tonic dopaminergic tone was thought encode motivation, with the fast phasic dopamine release encoding prediction-error learning (Schultz, 2007). However, recent work has challenged this ‘slow tonic = motivation’ and ‘fast phasic = learning’ idea of dopamine function. Dopamine release driven changes in motivation were found to occur as fast as the measurement techniques allows it to (i.e., ultra-fast (milliseconds) with dLight1, fast (seconds) with voltammetry and slow (minutes) with micro-dialysis) (Berke, 2018; Hamid et al., 2015; Mohebi et al., 2019). Therefore, dopamine-dependent changes in motivation can occur fast (sub-second scales), even within trials themselves, and do not necessarily conform to the slow tonic changes. The papers also found that rats were more motivated (i.e., they had shorter latency to start the next trial) when the average reward rate was high - when the animal is in a high dopaminergic state. Our current model has salience calculated at each state within the trial, allowing for fast modulation of the current action’s value. Also, dopamine levels ramp up as the animal gets closer to the reward (Howe et al., 2013), consistent with dopamine’s role in motivation/performance processes, interpreted as continuous increases in value states that are closest to the reward state (Berke, 2018). These dopamine ramps and the increasing value of states closest to the reward interpretation brings an interesting explanation into how relapse behaviors in people with substance use disorders increase as they get closer to the substance of abuse (high availability) (Abbott, 2020; Sureshkumar et al., 2021; Washton, 1986). Here, a higher drug availability or proximity to the drug may cause dopamine levels to ramp up, causing a high value representation for the states that are closest to the drug reward and increasing the probability of relapsing as a result. One consequence of these dopamine ramps is a higher average prediction-error rate, and this would create a salience misattribution effect where the positives of the current action are overweighted compared to the negatives (eq 8). Therefore, as the person gets closer to the drug, and dopamine ramps up, the value of that current action (taking the drug) is increased. This is because the positives of the current drug action are weighted even more, and at the same time, the negatives are weighted less. This causes an even larger value increase towards the drug action but slows down any value decreases towards that drug action. Therefore, dopamine encoding motivation/performance has an important implication for how relapse may be made more likely through the salience misattribution effect in eq 8.

One possibility for how learning and motivation may work together under the present model is through different stages within a trial. When a drug-cue, or any cue with a high salience is presented, it may trigger the motivational salience and drive the decision towards the states that follow the high salience cue, which is consistent with the *incentive salience* accounts (Robinson & Berridge, 1993). Data from Hamid et al., (2015) is also consistent with this cue triggering motivational salience interpretation, where artificial stimulation of dopaminergic neurons within the VTA (via optogenetics) caused the animal to engage faster (with more motivation) at the current trial (consistent with dopamine’s motivation/performance). But at the stage where a reward prediction-error is generated, it may be the learning/updating processes involved. This learning stage is where the prediction-errors are weighted in proportion to the salience placed on them (i.e., more from positive prediction-errors, and less from negative). This learning interpretation during the reward prediction error stage is also consistent with data from Hamid et al., (2015). Here, artificial dopaminergic stimulation during a reward prediction-error caused the animal to repeat the same action (consistent with learning/value update role of dopamine). That data can be interpreted under our salience model where the increased dopamine is now causing a greater value update of that same action (by over-weighting positives and downweighting negatives), and therefore increasing the probability of repeating the same action. This interpretation is particularly supported because the same action is repeated, irrespective of whether the reward was given or not at the end of the trial. Therefore, when the negatives of an action are downweighted in salience, and any positives are overweighted – the probability of repeating that action, regardless of the outcome, is made higher. Similarly, in the case of addictive behaviors, the agent is reinforced to take drug actions through cue triggered motivational salience (that accelerates the value increase by downweighting negatives and overweighting positives of that action). The agent is then further reinforced through asymmetric learning from the prediction-errors generated at the feedback stage, where positives are more heavily weighted, and negatives less so. This salience misattribution effect may be one motivation and learning dysfunction in people with a substance use disorder which could further increase drug-related behaviors. This idea is speculative but could be tested experimentally, wherein midbrain dopaminergic signatures (release and cell firing) are analyzed separately at the cue and prediction-error/feedback epochs of an experiment manipulating reward rate.

Given our strong focus on dopamine’s role in encoding reward prediction-error learning and motivation, it is critical to address recent experimental work suggesting that while dopamine encodes reward prediction-errors, it only does so in limited contexts/conditions (Coddington et al., 2023; Coddington & Dudman, 2018, 2019; Howe et al., 2013; Jeong et al., 2022; Kutlu et al., 2021; Matsumoto & Hikosaka, 2009; Ungless et al., 2004). Recently, Kutlu et al., (2021) suggested that dopamine’s role can be better explained as encoding *perceived salience*, instead of encoding reward prediction-errors and signed/unsigned non-reward prediction-errors. This brought together seemingly disparate results on dopamine’s role in prediction-errors and aversion. The paper used dLight1.1 to record dopamine dynamics with a fast (sub-second) temporal resolution *in vivo* in the nucleus accumbens core (NAc) in mice. Kultu et al., (2021) found that dopamine does encode reward prediction-errors during positive reinforcement (cue → action → reward) but not during negative reinforcement (cue → action → avoid punishment). Also inconsistent with the reward prediction-error hypothesis, when a cue predicts a reward, but then switches to now predicting a punishment (worse than expected), there was a positive dopamine response, instead of negative which the reward prediction-error hypothesis would predict. Overall, through various experiments, the authors suggested that dopamine is encoding perceived salience, which is computationally defined as the product of stimulus intensity and attentional value of a stimulus (termed the Kutlu-Calipari-Schmajuk (KCS) model). The attentional value factor is largely modulated by novelty, which is encoded by dopamine. Further, the authors note that perceived salience, particularly the attentional value factor, is subjective and depends on the context, situations, and experience and that it can contribute to learning.

In contrast to the KCS model, the SMMA model presented here views salience as being proportional to the instantaneous prediction-error and the average prediction-error. Kutlu et al., (2021) brought together seemingly disparate data in dopamine encoding reward prediction-errors and aversive stimuli under the perceived salience framework. However, our current model views salience as a direct modulator of learning and motivation within the reinforcement learning framework, which may not be inconsistent with the perceived salience framework. For example, after a switch from cue → reward, to now, cue → cost, our model would also predict a positive dopamine response, encoding salience. Here, a large negative prediction-error would increase the salience (eq 8), predicting a positive, and not a negative dopamine response. The difference in the SMMA model is that this salience is then used to update the value representation of that state, but this update is not directly conceptualized within the KCS model.

The SMMA model also predicts a positive salience-related dopamine response to novelty, through the average prediction-error term. The more the average prediction-error, the more novel the environment (i.e., if the agent has a high prediction-error rate, the environment is less known and produces large prediction-errors), and the greater the dopaminergic salience. The KCS model goes further to also explain dopamine’s role (or lack of a role) in non-reward related signed and unsigned prediction-errors, which our present salience model does not. Non-reward prediction-errors are not within the scope of the current model and is therefore a limitation, when compared to the KCS model. Both models are similar in that dopamine is encoding salience but diverge in how salience in mathematically defined and conceptually used to influence behavior. Our current model explains behavior through salience modulating value updating and explains aspects of asymmetric learning when in different dopaminergic states, whereas the KCS model deals with behavior change through novelty related attentional mechanisms and goes beyond reward prediction-errors.

The ACTR policy learning model by Coddington et al., (2023) is another new model of note in comparison to our model. The authors experimentally showed that optogenetic stimulation of midbrain dopaminergic neurons in rats could also slow learning, which is inconsistent with the traditional reward prediction error account of dopamine function. These findings were interpreted as mesolimbic dopamine activity playing a role in setting an adaptive learning rate of given action policies, which was a component within their ACTR model. Similar to our current model, the ACTR model has a dynamic parameter that weights learning on a trial-by-trial basis. However, within the ACTR model, the parameter modulating learning rate is defined based on the summation of sensory cue (preparatory) and movement/action (reactive) related components. This parameter is then used to weight performance-errors (and not reward prediction-errors) that drives policy learning. In the current model, learning was modulated through a salience factor based on the instantaneous and average reward prediction-errors, instead of updates in action policy values. In sum, similar to the other proposed models, we also propose that dopamine is playing a role in modulating learning. The critical distinction is that we propose this role for dopamine within the reward-prediction error framework, under a single parameter regime, while also integrating with the motivation function of dopamine.

### Anterior cingulate cortex and internal representation updates

We primarily focused on two factors in our salience model: the instantaneous prediction-error and the average prediction-error. While dopaminergic mechanisms are very likely involved in encoding these reward prediction-error processes, the average prediction-error may also be encoded in systems other than the dopaminergic system. Kennerley et al., (2006) report that ACC lesions in monkeys did not impair immediate performance in a reversal learning task but impaired the overall performance as the monkeys were less able to integrate overall reward histories to guide their choices. Wittmann et al., (2016) used functional magnetic resonance imaging (fMRI) in humans and found instantaneous prediction-error encoded in the ventral striatum blood oxygen level dependent (BOLD) signal. However, they also found the ACC encoded expected prediction-error, as calculated based on reward histories for a given environment. They hypothesized that expected prediction-error could modulate the weight placed on the instantaneous prediction-error.

Further accounts have also suggested that the ACC may encode reward histories and influence future behavior by integrating or weighting the immediate reward with the rate of rewards from previous trials (i.e., average reward rate) (Bernacchia et al., 2011; Buckley et al., 2009; Holroyd & Yeung, 2012; Sallet et al., 2007; Seo & Lee, 2007). For example, (Buckley et al., 2009) found that only a specific lesion in the macaque ACC, and not the other PFC regions, impaired performance in a Wisconsin-card sorting task such that reward histories were not integrated to optimize decisions. Other lesions, such as the orbitofrontal cortex, impaired the current reward value updating, but not the ACC-dependent use of reward histories. Another paper found that a lesion of the medial prefrontal cortex (mPFC) in rats, which includes the ACC, can disrupt weighting of reward prediction-errors within the ventral tegmental area (Starkweather et al., 2018). This shows that ACC might modulate instantaneous reward prediction-errors in the VTA. These studies collectively suggest that one possible way in which the salience in the SMMA model may be implemented is through ACC function, which may track the average prediction-error variable, accounting for reward histories in a given environment. This influences the instantaneous prediction-error and internal representation updating.

The involvement of ACC in estimating the average prediction-error may explain asymmetric learning in people with a substance use disorder through more than the increase in dopamine levels from the drug taking. Those with a substance use disorder may have a higher estimate for the average prediction-error due to reduced sensitivity/updating from error learning but increased sensitivity/updating from rewards. As a result, when the average prediction-error is calculated for people with a substance use disorder, the positives are represented more so than the negatives, possibly producing a higher average prediction-error. This high average prediction-error may then be used to further reduce error learning as proposed in our mathematical definition of salience, creating a cyclic process of decreased sensitivity to negatives but increased to positives. We speculate that this salience misattribution process may be initiated by dopamine from drugs or drug cues from the hypersensitized dopaminergic circuits, but then persists through the ACC weighting mechanism. We therefore predict that in an experiment where participants need to infer an average reward rate, people with a substance use disorder may have a higher estimation of this average reward rate than controls, particularly when drug cues are used to predict future rewards. In sum, both dopamine and ACC may play a role in weighting internal representation updating based on the average prediction-error and the instantaneous prediction-error (i.e., salience in eq 8) and producing the salience misattribution effect where positives are overweighted, compared to the negatives in people with a substance use disorder.

### Craving

Consistent with previous accounts (Ashare & Hawk, 2012; Giordano et al., 2002; Hoffman et al., 2008), our SMMA model produced steeper discounting for drug rewards under a craving state where the average prediction-error is negative, (Figure 12). However, the model also placed greater salience on non-drug related negative prediction-errors, compared to non-drug related positive prediction-errors under an induced craving state (where the average prediction-error is negative). The evidence for this effect for non-drug rewards during a craving state in people with a substance use disorder is currently mixed, with some suggesting reduced processing of both, positive and negative non-drug related reward prediction errors (Deserno et al., 2015; García-García et al., 2017; Park et al., 2010; Parvaz et al., 2015; Ubl et al., 2014). Rose et al., (2014) found an increased responsiveness to negative prediction-errors and decrease to positive in people with a cocaine use disorder. In contrast, Wang et al., (2019) found increased sensitivity to positive prediction errors and learning rates for loss avoidance in deprived people with a cocaine use disorder. Overall, our model predicts increased learning from negative prediction-errors and reduced learning from positives during the craving state. However, this prediction does not bring together the currently limited, but disparate data on cravings and prediction-errors, and is therefore a limitation of our current model.

There are other computational models of addiction that include craving but do so very conceptually. Redish et al., (2007) does not directly model craving but suggests that high drug availability and the presence of drug cues, may cause an agent to revert to the original state-representation, in which the drug actions were originally learned and where the value of drug rewards is very high. Reverting to this original state-representation would produce strong increases in craving levels and also the chances of relapsing. This model offers an explanation into why relapse may occur even after years of abstinence (Washton, 1986). Our current SMMA model does not include a state splitting component but may still explain aspects of cravings under one dynamic state, with changes in the average prediction-error.

Gu and colleagues used a Bayesian framework to conceptualize craving as a high ‘discomfort’ state due to failures in updating Bayesian beliefs on physiological/interoceptive bodily states (Gu, 2018; Gu & Filbey, 2017). Here, agents start with a prior belief of low craving/discomfort, as the drug is expected to be delivered, but then receive sensory evidence (Bayesian likelihood) associated with no drugs being delivered (e.g., increased heart rate). Over time, this sensory evidence starts to have high Bayesian precision as drugs continue to not be delivered. Hence, the agent becomes more certain of the lack of drug delivery, leading to an update in the posterior Bayesian belief towards increased discomfort/craving. Data from people with a nicotine use disorder suggests that this craving/discomfort state only reduces when the drug is expected (prior is towards low craving) and delivered, but not when the drug is unexpected but still delivered (Gu et al., 2016). Therefore, the model and the associated data from Gu et al., (2016) suggests that craving has a top-down component beyond just pharmacology, and that expectations on receiving the drug can influence whether there is a reduction in the craving state.

Both models (Gu, 2018; Redish et al., 2007) offer largely conceptual explanations of craving behaviors, but do not directly simulate craving behaviors, particularly behaviors on how drug and non-drug rewards may be differently processed under this craving state. Our current model views craving as a state where there is a negative average prediction-error, which causes more value updates from non-drug negatives than positives. This is more consistent with a depressive state (Rouhani & Niv, 2019) and may also be conceptualized as a ‘discomfort’ state, consistent with Gu & Filbey (2017). However, SMMA offers a more mathematically concrete conceptualization. Further, our model can simulate steeper discounting of drug rewards, which previous models do not. An important distinction of the proposed model is that the craving state can be induced by first giving the agent the drug and forming the expectation, and then removing the drug. This causes large negative prediction-errors to the point where the average prediction-error is negative. However, more human data and mathematically concrete models are needed in order to better understand the diverse neurocomputational underpinnings of the craving state.

### Delay Discounting

Delay discounting of rewards is a large component of the current model and is produced through placing less salience on the delay as the average prediction-error rate increases. Note that all the delay discounting effects here are produced without manipulating the discounting factor (γ) factor in eq 8, and thus are a consequence of a salience misattribution effect. Steeper discounting for delayed rewards is a complex phenomenon and involves many different neural systems and computations (Madden & Bickel, 2010). Pine et al., (2010) suggested that high dopamine levels, induced through L-dopa consumption, increases steepness in delay discounting in humans. However, dopamine’s role in modulating delay discounting is complex. There are mixed results based on the animal species used, the task and technique, the dose, and the drug used to modulate dopamine levels (Floresco et al., 2008; Isles et al., 2003; Kobayashi & Schultz, 2008; Koffarnus et al., 2011; Richards et al., 1999; Tedford et al., 2015; Wade et al., 2000). We limit our interpretations here to cases where dopamine may produce steeper discounting, with a high average prediction-error rate as a consequence of increased dopamine levels. This is in line with Niv et al., (2007), who suggest that a high reward rate causes the agent to act with more vigor and, with less engagement of the deliberative decision-making strategies, as the opportunity cost of not acting in a rich environment is greater than the cost of acting. We propose that one consequence of these high vigor behaviors in a rich environment, with a high average reward and prediction-error rate, is that the delay has less salience, producing less value updating for delayed rewards, which may therefore contribute to faster discounting.

In a task involving episodic future thinking and delay discounting, using fMRI in humans, Peters & Büchel (2010) found increased engagement of the ACC and ACC-hippocampal coupling when participants were reminded of the future event compared to when they were not. The authors interpreted this as the ACC engaging greater cognitive control and adjusting values based on the changes in context, supporting adaptive decision-making. We speculate that ACC engagement may influence delay discounting by increasing the salience of future events, possibly through initiating hippocampal task representations of the future events, making the future more concrete and producing slower discounting. Previous accounts have found that a mPFC disruption reduced vicarious trial and error (VTE) behaviors (Kidder et al., 2021; Schmidt & Redish, 2021). VTE is when an animal imagines the future by initiating a representation of future outcomes for evaluation and deliberates one choice over the other (Johnson & Redish, 2007; Redish, 2016). A neurophysiological basis for these VTE task representations are within the hippocampal place cells (Johnson & Redish, 2007). Given that mPFC disruption reduces these deliberative VTE behaviors, we speculate that ACC engagement here may be initiating the task representation though hippocampal coupling, such that the future is represented with more salience, and increasing the value update, and decreasing the discounting of the future as the delay increases. Overall, states with high dopamine levels, where there is a high average prediction-error, may cause behaviors with increased vigor, where ACC-hippocampal dependent deliberate decision-making strategies are less engaged, and may therefore produce steeper discounting in people with a substance use disorder whereby the future may be represented with less certainty, and therefore less salience.

### Behavioural addictions

It has been suggested that the environmental contingencies of some situations could produce behavioural addictions through continuous positive prediction-errors (e.g., endless scrolling for social media and perpetual reward uncertainty for gambling) (Ciria et al., 2022; Zack et al., 2020). Further, high unpredictability and uncertainty may be used by designers to possibly increase use and maximise positive prediction-errors through unexpected positive notifications (e.g., more “likes” than expected) (Alter, 2017; Lanier, 2019). The reversal learning simulation in Figure 8, without the ncDA factor, suggests that even a high average prediction-error rate, which is caused by a large number of positive prediction-errors, can produce a salience misattribution effect where the instantaneous positive prediction-errors are weighted more so than the negatives. As a result, in a positive but unpredictable environment, the agent is more likely to win (get a positive prediction-error) and stay as a result. Likewise, the agent is less likely to lose (get a negative prediction-error) and change behaviour. Therefore, unpredictability can enhance behavioural addictions in part because of this salience misattribution effect, causing the agent to be less likely to switch behaviour even after losses/negative prediction-errors and more likely to win and stay on the same behaviour. The current model predicts that the point where the agent is most likely to stay in the addictive behaviour is when the ratio between the absolute prediction-error and the average prediction-error is close to 1. It is at this point where the difference between the salience placed on the negative prediction-error and the positive prediction-error is the greatest (see Fig. 8d), making the agent least likely to lose and shift behaviour and most likely to win and stay on the same behaviour. The decrease in lose-shift effect is bounded by the absolute ratio of the average prediction-error and the prediction-error to be less than 1, and when the ratio is above 1, lose-shift starts to increase. This increase in lose-shift behaviour is possibly a limitation of the current model, as a high positive prediction-error rate should continue to make the agent less likely to lose and shift, and this limitation should be addressed in future models that are more specifically designed to address behavioural addictions.

### Conclusions and future directions

Starting from the concept that salience modulates internal representation updating, we proposed a novel mathematical model where salience is proportional to the instantaneous prediction-error and the average prediction-error, and the salience placed on the delay to the reward is inversely proportional to the average prediction-error rate. Using this definition, we simulated key aspects of addictive behaviours where 1) drug related negative prediction-errors/costs are downweighted in salience, but 2) drug related positive prediction-errors/rewards are overweighted in salience, and 3) drug rewards are discounted faster than non-drug rewards. We also conceptualized a craving statein the model as a state with a negative average prediction error where drug rewards are discounted faster, and non-drug positives are weighted less than non-drug negatives. We went further to suggest that maximising positive prediction-errors and increasing the average prediction-error rate may contribute to the behavioural persistence seen in behavioural addictions (e.g., social media and gambling). Here, a high average prediction-error contributes to the salience misattribution effect where the costs are downweighted in salience, and consequently, the probability of losing and shifting is reduced. At the same time, rewards are overweighted in salience and consequently, the probability of winning and repeating the same behaviour increases. In sum, we suggest that a salience factor that modulates internal representation updating could be producing a misaligned internal representation in people with a substance use disorder. This misaligned internal representation is then used to produce key aspects of maladaptive decision-making in people with a substance use disorder. The proposed model may be relevant in two main future research directions: 1) experimentally testing how dopamine and ACC may regulate reward-related motivation and learning through a salience factor that modulates internal representation updating and, 2) how this salience misattribution effect may be ameliorated in people with a dependence, so that the internal representation is no longer misaligned in selectively updating from drug related positives, and less so from drug related costs.

## Notes

### Competing Interest Statement

The authors have declared no competing interest.

