## Supplementary Materials for "Reward prediction-errors weighted by cue salience produces addictive behaviors in simulations, with asymmetrical learning and steeper delay discounting"

### Methods

*Increased salience on drug cues*

A 3-state space model was used here where agents that go from S1→S2→ S3, which is 1 trial, and then back to S1 for the next trial (see the state-space in Figure 1). There were a total of 300 trials in this simulation. When simulating using the SMMA and the Redish 2004 model, agents received either a drug reward (r = 5 and ncDA = 0.5) or a non-drug reward (r = 5) in state 3, with state 2 being the cue state and the value and prediction-errors plotted from this state. In the SMMA model, the changing salience was applied to the cue state (state 2). When simulating using the Zhang et al. 2009 model, reward was delivered in state 3, with the changing salience applied in state 3, and the value/prediction-errors were plotted from this state.

*Lever presses for drug rewards*

The state-space simulating lever presses for drug rewards using the current model is in Figure 2a and Zhang et al., (2009) model in Figure 2b. In both models, agents decide between action *a1* for a non-drug reward or action *a2* for a drug reward. The salience for the non-drug reward is 1, with the salience varied for the drug-reward, applied at the cue state for the present model and at the reward state for Zhang et al., (2009) model. There were a total of 1000 timesteps, with 1 trial/decision being 3 timesteps. Salience was varied from 1 to 8.25, with incremental an increase of 0.25. The same procedure was followed for the Redish 2004 model, except there was no salience factor.

*Modelling probability of taking drug actions given contrasting non-drug rewards*

The state-space model used for the present simulation is on Figure 3, for both the Redish 2004, the SMMA model and the Zhang et al. 2009 model. Here, agents made decisions between the drug reward (*a1; r = 5 and ncDA = 0.5 (for both models)*) or the non-drug reward (*a2*) which was varied between 1 and 25, with increments of 0.25 for each simulation. There were a total of 2000 trials/decisions made by the agent. The proportion drug actions were calculated as the number of drug actions taken divided by the total number of actions taken. The proportion of drug actions were calculated in increments of 250 trials, up to 1000 actions (e.g., proportion of drug actions taken in the first 250 trials).

*Developing inelasticity over time*

The state-space model used for the simulation is on Figure 4, for both the Redish 2004, the SMMA and the Zhang et al. 2009 models. Here, agents chose between action *a1* which is to stay in the current state and avoid the cost but also avoid the drug reward or take action *a2* which comes with a given cost, but also with a drug reward. There was a total of 10,000 timesteps in each simulation. Cost was varied between 0 and 4, with increments of 0.25 for each simulation. The probability of taking the drug action calculated in increments of 100 drug actions, i.e., for the first 100 drug actions taken (100 divided the total number of timesteps to make 100 drug actions), then the next 100 drug actions (100 divided by the total number of timesteps taken to make the next 100 drug actions), up to 400 drug actions.

*Salience attribution on negative and positive prediction-errors using the current SMMA model*

The state-space model used for the present simulation is on Figure 5. The agents received a non-drug reward of 5 in state 3 and another non-drug reward of 5, in state 5. After 300 trials, the agent received a reward of 7 in state 3 (causing a positive prediction-error of 2). This simulation was repeated with the agents now receiving a cost of 5 in state 3, and then a cost of 7 after 300 trials in the same state (causing a negative prediction-error of 2). Lastly, both these simulations were repeated with the reward being a drug reward in state 5 (reward = 5, ncDA = 0.5). All these simulations ran for 330 trials in total, with the positive or negative prediction-error given at trial 300.

*Reversal learning with drug and non-drug rewards*

There were two different types of simulations performed here – one with the drug reward in state 6 and another with no drug reward, in any of the states (see Figure 7a-b). In the simulation with no drug rewards (Figure 7b), agents chose between action *a1* which led to a reward in state 2 (r = 1) followed by another reward in state 6 (r = 5) or action *a2* where the agent would receive a cost (r = -1) at state 3, followed by another reward at state 7 (equal to the reward in state 6, r =5). Therefore, action *a1* would be a win, and action *a2*, a loss. After every 100 trials, there was a reversal where the cost would be given for taking action *a1* (in state 2) and reward for action *a2* (in state 3). There were a total of 1000 trials, with a reversal of this type after every 100 trials. Simulation two is identical to this, except the reward given at state 6 would be a drug reward (Figure 7a). The inverse temperature parameter used here was 0.25 instead of 0.5 which was used in all other simulations. This was done to demonstrate the differences between drug-related and non-drug related actions, without reaching ceiling levels for the two simulations. Probability of win-stay was calculated as the proportion of actions repeated right after the trial where reward was received in state 2 or 3 (number of win-stay trials divided by all the win trials). Probability of lose-shift was calculated as the proportion of actions not repeated following a cost at state 2 or 3 (number of lose-shift trials divided by all the lose trials).

*Reversal learning with non-drug rewards and variable average prediction-errors; the link with behavioral addictions*

Here, agents received a reward of 5 for taking action *a1* or a cost of 5 for taking action *a2* (see Figure 8). There were a total of 1000 trials, with a reversal every 100 trials. In this reversal, there would now be a cost of 5 for talking action *a1* and a reward of 5 for taking action *a2*. There was a mean prediction-error made to vary between 0 and 5.9, with increments of 0.1 per simulation. A win was defined as taking the action with the reward, and loss defined as taking the action that led to the cost. Probability of win-stay was calculated as the proportion of actions repeated right after the trial where reward was received in state 2 or 3 (number of win-stay trials divided by all the win trials). Probability of lose-shift was calculated as the proportion of actions not repeated following a cost at state 2 or 3 (number of lose-shift trials divided by all the lose trials).

*Delay discounting simulations*

All delay discounting simulations followed a 3-state model, where the variable delay was given in state 2, with the reward received in state 3. There were a total of 300 trials in all these simulations, with the value taken from state 2 (where the delay was given), after the 300 trials. The values were normalized between 0 and 1 to compare between simulations. The first delay discounting simulation involved discounting drug and non-drug rewards. These were carried out separately, first with the non-drug reward of 5. Following this a separate simulation was carried out with the drug reward (r = 5 and ncDA = 0.5) (Figure 9). Next, simulation was carried out with a variable mean prediction-error for drug rewards (see Figure 10). This was identical to the previous simulation with the drug reward except now the mean prediction-error parameter was varied, with values of 0,0.5,1 and 2.5. Each of these mean prediction-error values comprised of a separate simulation. Lastly, the craving simulation (Figure 12) was carried out the same except now the mean prediction-error value was -2.5.

*Induced craving, bingeing, and salience simulations*

Craving was induced through two steps. The first step involved generating the expectation of receiving the drug reward (for 30 trials). Following this, the drug reward was no longer given, causing negative prediction-errors to the point where the mean prediction-error became negative (by trial 75; 45 trials after drugs were removed). At this point (when mean prediction-error was negative), either a negative or a positive prediction-error (of 1.75) was given, and the salience placed on these negative and positive-prediction errors were plotted (see Figure 11). The bingeing state was given the same prediction-error but with the drug rewards never being removed. There were a total of 100 trials in this simulation.
